# Efficient and Robust Search of Microbial Genomes via Phylogenetic Compression

**DOI:** 10.1101/2023.04.15.536996

**Authors:** Karel Břinda, Leandro Lima, Simone Pignotti, Natalia Quinones-Olvera, Kamil Salikhov, Rayan Chikhi, Gregory Kucherov, Zamin Iqbal, Michael Baym

## Abstract

Comprehensive collections approaching millions of sequenced genomes have become central information sources in the life sciences. However, the rapid growth of these collections has made it effectively impossible to search these data using tools such as BLAST and its successors. Here, we present a technique called phylogenetic compression, which uses evolutionary history to guide compression and efficiently search large collections of microbial genomes using existing algorithms and data structures. We show that, when applied to modern diverse collections approaching millions of genomes, lossless phylogenetic compression improves the compression ratios of assemblies, de Bruijn graphs, and *k*-mer indexes by one to two orders of magnitude. Additionally, we develop a pipeline for a BLAST-like search over these phylogeny-compressed reference data, and demonstrate it can align genes, plasmids, or entire sequencing experiments against all sequenced bacteria until 2019 on ordinary desktop computers within a few hours. Phylogenetic compression has broad applications in computational biology and may provide a fundamental design principle for future genomics infrastructure.

## INTRODUCTION

Comprehensive collections of genomes have become an invaluable resource for research across life sciences. However, their exponential growth, exceeding improvements in computation, makes their storage, distribution, and analysis increasingly cumbersome ^1^. As a consequence, traditional search approaches, such as the Basic Local Alignment Search Tool (BLAST) ^2^ and its successors, are becoming less effective with the available reference data, which poses a major challenge for organizations such as the National Center for Biotechnology Information (NCBI) or European Bioinformatics Institute (EBI) in maintaining the searchability of their repositories.

The key to achieving search scalability are compressive approaches that aim to store and analyze genomes directly in the compressed domain ^3,4^. Genomic data have low fractal dimension and entropy ^5^, offering the possibility of efficient search algorithms ^5^. However, despite the progress in compression-related areas of computer science ^4–15^, it remains a practical challenge to compute parsimonious compressed representations of the exponentially growing public genome collections.

Microbial collections are particularly difficult to compress due to the huge number of genomes and their exceptional levels of genetic diversity, which reflect the billions of years of evolution across the domain. Even though substantial efforts have been made to construct comprehensive collections of all sequenced microbial genomes, such as the 661k assembly collection ^16^ (661k pre-2019 bacteria) and the BIGSIdata de Bruijn graph collection ^17^ (448k de Bruijn graphs of all pre-2017 bacterial and viral raw sequence), the resulting data archives and indexes range from hundreds of gigabytes (661k) to tens of terabytes (BIGSIdata). This scale exceeds the bandwidth, storage, and data processing capacities of most users, making local computation on these data functionally impossible.

We reasoned that the redundancies among microbial genomes are efficiently predictable, as they reflect underlying processes that created the collection: evolution and sampling. While genomes in nature can accumulate substantial diversity through vertical and horizontal mutational processes, this process is functionally sparse, and at the same time subjected to selective pressures and drift that limit their overall entropy. The amount of sequenced diversity is further limited by selective biases due to culture and research or clinical interests, resulting in sequencing efforts being predominantly focused on narrow subparts of the tree of life, associated with model organisms and human pathogens ^16^. Importantly, such subtrees have been shown to be efficiently compressible when considered in isolation, as low-diversity groups of oversampled phylogenetically related genomes, such as isolates of the same species under epidemiological surveillance ^18,19^. This suggests that the compression of comprehensive collections could be informed by their evolutionary history, reducing the complex problem of general genome compression to the more tractable problem of local compression of phylogenetically grouped and ordered genomes.

Phylogenetic relatedness is effective at estimating the similarity and compressibility of microbial genomes and their data representations. The closer two genomes are phylogenetically, the closer they are likely to be in terms of mathematical similarity measures, such as the edit distance or *k*-mer distances ^20^, and thus also more compressible. Importantly, this principle holds not only for genomes, but also for de Bruijn graphs and many *k*-mer indexes. We reasoned that phylogenetic trees could be embedded into computational schemes in order to group similar data together, as a preprocessing step for boosting local compressibility of data. The well-known Burrows-Wheeler Transform ^21^ has a similar purpose in a different context and similar ideas have been used for read and alignment compression ^22–25^. Other related ideas have previously been used for scaling up metagenomic classification using taxonomic trees ^26–29^ and search in protein databases ^30,31^.

At present, the public version of BLAST is frequently used to identify the species of a given sequence by comparing it to exemplars, but it is impossible to align against *all* sequenced bacteria. Despite the increasing number of bacterial assemblies available in the NCBI repositories, the searchable fraction of bacteria is exponentially decreasing over time (**Fig. 1a**). This limits our ability to study bacteria in the context of their known diversity, as the gene content of different strains can vary substantially, and important hits can be missed due to the database being unrepresentative.

**Fig. 1:**
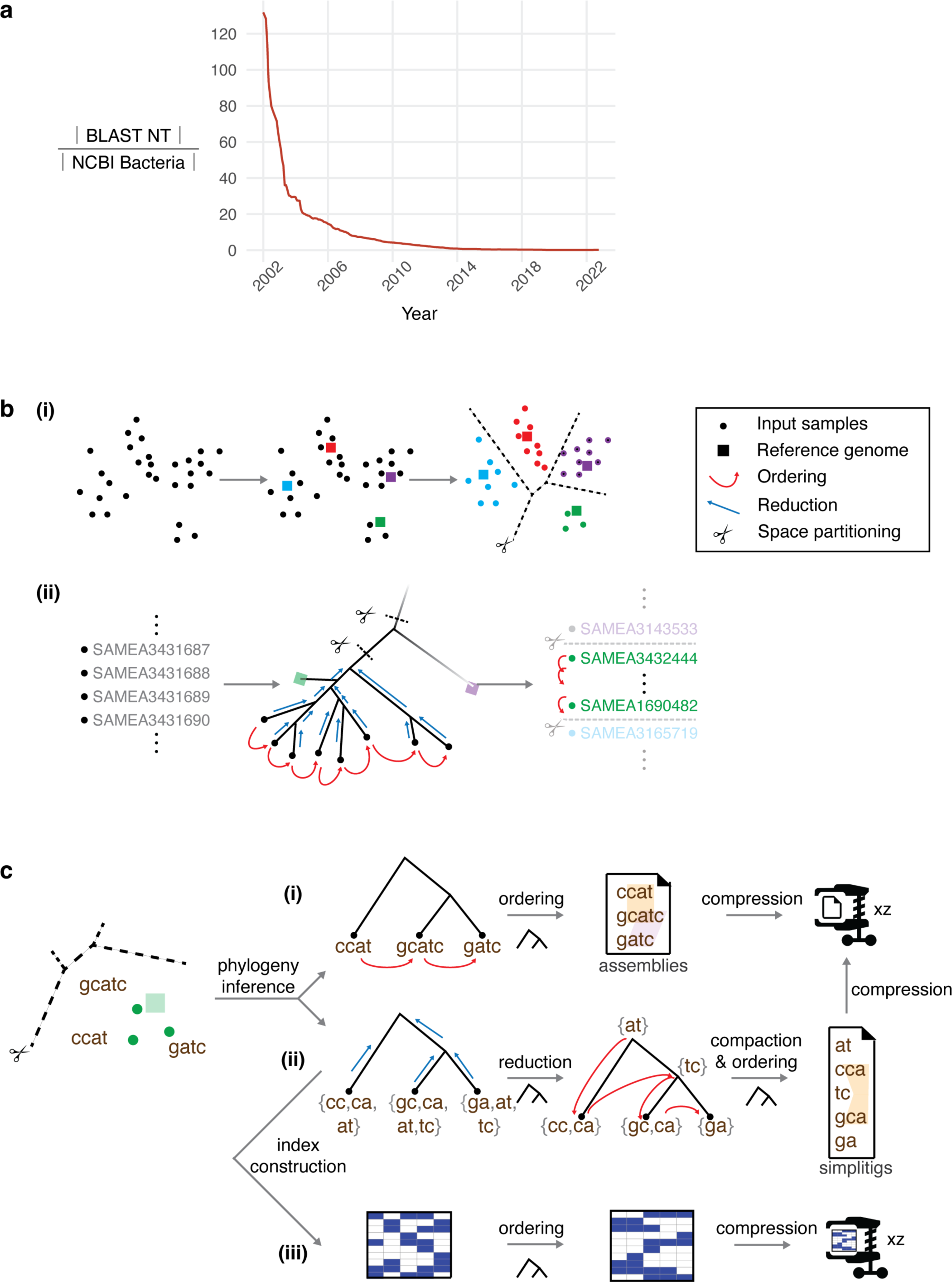
Overview of phylogenetic compression and its applications to different data types. **a)** Exponential decrease of data searchability over the past two decades illustrated by the size of the BLAST NT database divided by the size of the NCBI Bacterial Assembly database. **b)** The first three stages of phylogenetic compression prior to the application of a low-level compressor/indexer. **(i)** A given collection is partitioned into size-and diversity-balanced batches of phylogenetically related genomes (e.g., using metagenomic classification of the original reads). **(ii)** The input data are reversibly reordered based on a compressive phylogeny, performed separately for each batch. **c)** Examples of specific protocols for phylogenetic compression of individual data types, performed separately for each batch. **(i)** Assemblies are sorted left-to-right according to the topology of the phylogeny, and then compressed using a low-level compressor such as XZ ^7,32^ or MBGC ^18^. **(ii)** For de Bruijn graphs, *k*-mers are propagated bottom-up along the phylogeny, and the resulting *k*-mer sets are compacted into simplitigs ^33^, which are then compressed using XZ. **(iii)** For BIGSI *k*-mer indexes, Bloom filters (in columns) are ordered left-to-right according to the phylogeny, and then compressed using XZ.

Here, we present a solution to the problem of searching vast libraries of microbial genomes: *phylogenetic compression*, a technique for an evolutionary-guided compression of arbitrarily sized genome collections. We show that the underlying evolutionary structure of microbes can be efficiently approximated and used as a guide for existing compression and indexing tools. Phylogenetic compression can then be applied to collections of assemblies, de Bruijn graphs, and *k*-mer indexes, and run in parallel for efficient processing. The resulting compression yields benefits ranging from a quicker download (reducing Internet bandwidth and storage costs), to efficient search on personal computers. We show this by implementing BLAST-like search on all sequenced pre-2019 bacterial isolates, which allow us to align genes, plasmids, and sequencing reads on an ordinary laptop or desktop computer within a few hours, a task that was completely infeasible with previous techniques.

## RESULTS

We developed a technique called phylogenetic compression for evolutionarily informed compression and search of microbial collections (**Fig. 1**, https://brinda.eu/mof). Phylogenetic compression combines four ingredients (**Fig. 1b**): 1) *clustering* of samples into phylogenetically related groups, followed by 2) inference of a *compressive phylogeny* that acts as a template for 3) *data reordering*, prior to 4) the application of a calibrated *low-level compressor/indexer* (Methods). This general scheme can be instantiated to individual protocols for various data types as we show in **Fig. 1c**; for instance, a set of bacterial assemblies can be phylogenetically compressed by XZ (the Lempel-Ziv Markov-Chain Algorithm ^7^, implemented in XZ Utils ^32^) by a left-to-right enumeration of the assemblies, with respect to the topology of their compressive phylogeny obtained via sketching ^34^.

We implemented phylogenetic compression protocols for assemblies, for de Bruijn graphs, and for *k*-mer indexes in a tool called MiniPhy (Minimization via Phylogenetic compression, https://github.com/karel-brinda/miniphy). To cluster input genomes, MiniPhy builds upon the empirical observation that microbial genomes in public repositories tend to form clusters corresponding to individual species ^35^, and species for individual genomes can be identified rapidly via metagenomic classification ^36^ (**Fig. 1b**, Methods). As some of the resulting clusters may be too large or too small, and thus unbalancing downstream parallelization, it further redistributes the clustered genomes into size-and diversity-balanced batches (Methods, **Supplementary Fig. 1**). This batching enables compression and search in a constant time (using one node per batch on a cluster) or linear time (using a single machine) (Methods). For every batch, a compressive phylogeny – either provided by the user or computed automatically using Mashtree ^34^ / Attotree (https://github.com/karel-brinda/attotree, Methods) – is then used for data reordering (Methods). Finally, the obtained reordered data are compressed per batch using XZ with particularly optimized parameters (Methods), and possibly further re-compressed or indexed using some general or specialized low-level tool, such as MBGC ^18^ or COBS ^37^ (Methods).

We evaluated phylogenetic compression using five microbial collections, selected as representatives of the compression-related tradeoffs between characteristics including data quality, genetic diversity, genome size, and collection size (GISP, NCTC3k, SC2, 661k, and BIGSIdata; Methods, **Supplementary Table 1**). We quantified the distribution of their underlying phylogenetic signal (Methods, **Supplementary Table 2, Supplementary Fig. 2**), used them to calibrate the individual steps of the phylogenetic compression workflow (Methods, **Supplementary Fig. 3–5**), and evaluated the resulting performance, tradeoffs, and extremal characteristics (Methods, **Supplementary Table 3, Supplementary Fig. 6**). As one extreme, we found that 591k SARS-CoV-2 genomes can be phylogenetically compressed using XZ to only 18.1 bytes/genome (Methods, **Supplementary Table 3**, **Supplementary Fig. 4, 6**), resulting in a file size of 10.7 Mb (13.2× more compressed than GZip). A summary detailing the sensitivity/stability of performance to various factors is provided in **Supplementary Note 1**.

We found that phylogenetic compression improved the compression of genome assembly collections that comprise hundreds of thousands of isolates of over 1,000 species by more than an order of magnitude compared to the state-of-the-art (**Fig. 2a**, **Supplementary Table 3**). Specialized high-efficiency compressors such as MBGC ^18^ are not directly applicable to highly diverse collections, therefore, the compression protocols deployed in practice for extremely large and diverse collections are still based on the standard GZip, such as the 661k collection, containing all bacteria pre-2019 from ENA ^16^ (n=661,405, 805 GB). Here, MiniPhy recompressed the collection to 29.0 GB (27.8× improvement; 43.8 KB/genome, 0.0898 bits/bp, 5.23 bits/distinct *k*-mer) using XZ as a low-level tool, and further to 20.7 GB (38.9× improvement; 31.3 KB/genome, 0.0642 bits/bp, 3.74 bits/distinct *k*-mer) when combined with MBGC ^18^ that also accounts for reverse complements (**Fig. 2a**, **Supplementary Table 3**, Methods). Additionally, we found that the lexicographically ordered ENA datasets, as being partially phylogenetically ordered, can serve as an approximation of phylogenetic compression, with compression performance only degraded by a factor of 4.17 compared to full phylogenetic compression (**Supplementary Table 3**, Methods). The resulting compressed files are provided for download from Zenodo (**Supplementary Table 4**).

**Fig. 2:**
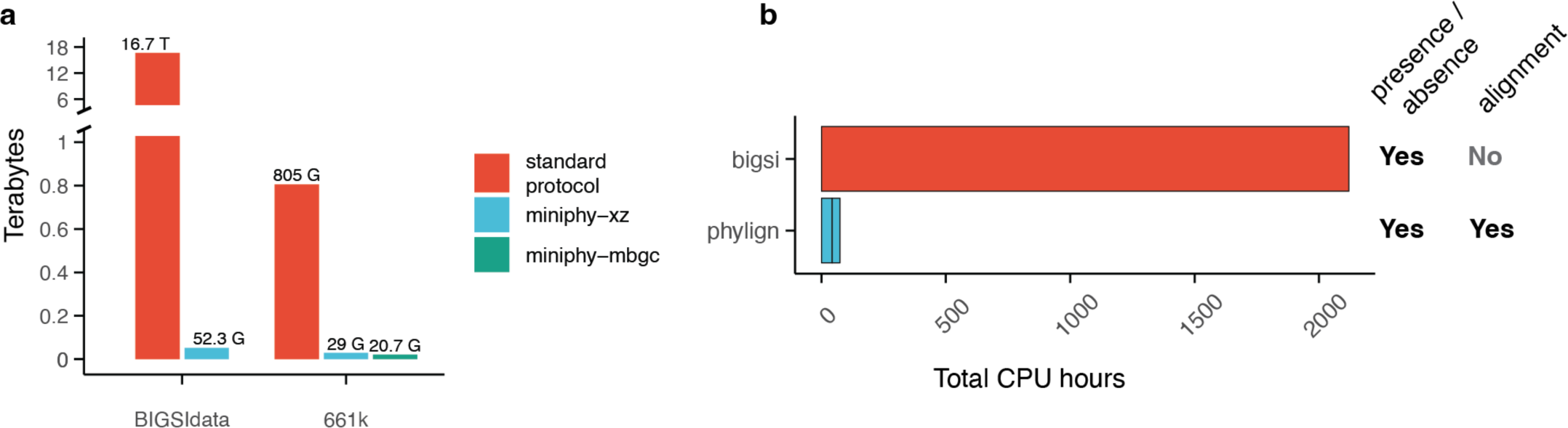
Results of phylogenetic compression. **a)** Compression by MiniPhy of the two comprehensive genome collections: BIGSI (425k de Bruijn graphs; the standard compression is based on McCortex binary files) and 661k (661k bacterial assemblies; the standard protocol is based on GZip). For BIGSIdata, MBGC is not included as it does not support simplitigs. **b)** Comparison of the Phylign vs. BIGSI methods on search of all plasmids from the EBI database. For Phylign, the two segments correspond to the times of matching and alignment, respectively.

We then studied de Bruijn graphs, a common genome representation directly applicable to raw-read data ^17,38^, and found that phylogenetic compression can improve state-of-the-art approaches by one-to-two orders of magnitude (**Fig. 2a**, **Supplementary Table 3,** Methods). As standard and colored de Bruijn graphs lack methods for joint compression at the scale of millions of genomes and thousands of species, single graphs are often distributed individually ^39^. For instance, the graphs of the BIGSIdata collection ^17^, comprising all viral and bacterial genomes from pre-2017 ENA (n=447,833), are provided in an online repository in the McCortex binary format ^40^ and occupy in total >16.7 TB (Methods). Here, we retrieved n=425,160 graphs from the Internet (94.5% of the original count) (Methods) and losslessly recompressed them using the MiniPhy methodology, with a bottom-up propagation of the *k*-mer content, to 52.3 GB (319× improvement; 123. KB/genome, 0.248 bits/unitig bp, 10.2 bits/distinct *k*-mer) (**Fig. 2a**, **Supplementary Table 3**, Methods). Further, as recent advances in de Bruijn graph indexing ^15^ may lead to more efficient storage protocols in the future, we also compared MiniPhy to MetaGraph ^38^, an optimized tool for indexing on high-performance servers with a large amount of memory. Here, we found that MiniPhy still provided an improvement of a factor of 5.78 (Methods).

Phylogenetic compression can be applied to any genomic data structure based on a genome-similarity-preserving representation (Methods, **Supplementary Note 2**). We demonstrate this using the Bitsliced Genomic Signature Index (BIGSI) ^17^ (**Fig. 1c(iii)**), a *k*-mer indexing method using an array of Bloom filters, which is widely used for large-scale genotyping and presence/absence queries of genomic elements ^16,17^. Using the same data, batches, and orders as inferred previously, we phylogenetically compressed the BIGSI indexes of the 661k collection, computed using a modified version of COBS ^37^ (**Supplementary Table 5**, Methods). Phylogenetic compression provided an 8.51× overall improvement compared to the original index (from 937 GB to 110 GB), making it finally usable on ordinary computers. After we further omitted the 3.7% genomes that had not passed quality control in the original study ^16^ (the 661k-HQ collection, visualized in **Supplementary Fig. 7**), the resulting phylogenetic compression ratio improved to 12.3× (72.8 GB) (**Supplementary Table 5**).

To better understand the impact of phylogenetic compression across the tree of life, we analyzed the 661k MiniPhy batches of assemblies and COBS indexes, both before and after compression (**Supplementary Fig. 8**). We found that although the top ten species constituted 80% of the genomic content, they occupied less than half of the database space post-compression for both genome representations (**Supplementary Fig. 8**). Conversely, the ‘dustbin’ batches, which include genomes from sparsely sampled species, expanded to occupy a proportion that was 9.4× larger in the database post-compression, compared to their precompression proportion, again for both representations (**Supplementary Fig. 8**). This consistent effect of compression on both assemblies and COBS indexes suggests that phylogenetic compressibility adheres to the same principles, irrespective of the specific genome representation used, with divergent genomes being a major driver of the final size.

To demonstrate the practical utility of phylogenetic compression, we used it to implement BLAST-like search across all high-quality pre-2019 bacteria for standard desktop and laptop computers (Phylign, http://github.com/karel-brinda/phylign, Methods). For a given a set of queries, Phylign first identifies for each query those genomes that match best globally across the whole 661k-HQ collection, by proceeding via progressive in-memory decompression and querying of individual phylogenetically compressed COBS ^37^ *k*-mer indexes (described above). Subsequently, Phylign iterates over the phylogenetically compressed genome assemblies (described above) and computes the corresponding full alignments using on-the-fly instances of Minimap 2 ^41^ (Methods). The choice of tools was arbitrary, and other programs or core data structures could readily be used instead. The resulting requirements amount to only 102 GB disk (for the compressed COBS indexes and assemblies: 195 KB/genome, 0.329 bits/bp, 23.0 bits/distinct *k*-mer) (**Supplementary Table 6**) and 12 GB RAM, and Phylign can thus be deployed on most modern laptop and desktop computers.

We first evaluated Phylign with 661k-HQ using three different types of queries – resistance genes (the entire ARG-ANNOT database of resistance genes ^42^, n=1,856), plasmids (EBI plasmid database, n=2,826), and a nanopore sequencing experiment (n=158,583 reads), with results available within 3.9, 11, and 4.3 hours, respectively, on an iMac desktop (**Supplementary Table 7**). Benchmarking against other tools was not possible, as we were unable to find any tool capable of aligning queries to 661k-HQ in a comparable setup. We therefore used the EBI plasmid dataset to compare Phylign to BIGSI with its original database of 448k genomes (which is essentially a subset of 661k-HQ with 1.43× less genomes) ^17^. We found that Phylign was over an order of magnitude faster (**Fig. 2b, Supplementary Table 7**); the search required 74.1 CPU hours and improved performance by a factor of 28.6× compared to the same BIGSI benchmark with its smaller database (**Fig. 2b, Supplementary Table 7**), while providing the full alignments rather than presence/absence only (**Fig. 2b)**. To our knowledge, this is the first time that alignment to a collection of a comparable size and diversity has been locally performed.

## DISCUSSION

It is hard to overstate the impact on bioinformatics of BLAST ^2^, which has allowed biologists across the world to simply and rapidly compare their sequence of interest with essentially all known genomes – to the extent that the tool name has become a verb. The web version provided by NCBI/EBI is so standard that it is easy to overlook how representative or complete its database is. However, twenty-four years on, sequencing data is far outstripping BLAST’s ability to keep up. Much work has gone into approximate solutions ^15^, but full alignment to the complete corpus of bacterial genomes has remained impossible. We have addressed this problem and made significant progress, via phylogenetic compression, a highly efficient general technique using evolutionary history of microbes to improve existing compressive data structures and search algorithms by orders of magnitude. More concretely, BLAST-like search of all microbes is now possible, not just for NCBI/EBI, but for anyone on a personal laptop. This has wide-ranging benefits, from an easy and rapid download of large and diverse genome collections, to reductions in bandwidth requirements, transmission/storage costs and computational time.

Elements of our approach and related techniques have been previously used in other contexts. Reversible reordering to improve compression forms the core of the Burrows-Wheeler Transform ^21^ and its associated indexes ^43–45^, and it has also been used for read compression ^22–25^. Tree hierarchies have been applied in metagenomics for both lossy ^26,27,46^ and lossless ^28^ reference data compression. Finally, a divide-and-conquer methodology has been employed to accelerate the inference of species trees ^47^. However, this is the first time all these ideas have been combined together to improve the scalability of search in large genome databases.

As with all forms of compression, our ability to reduce data is fundamentally limited by the underlying entropy. For genome collections, this is not introduced solely by the underlying genetic signal, but it is also tightly connected with the sequencing process and our capacity to reconstruct genomes from sequencing reads. The noise in the underlying *k*-mer histograms (**Supplementary Fig. 7**) suggests that any method for compression or search will have to address noise in the forms of contamination, missing regions, and technological artifacts, with legacy data posing a major challenge for both storage and analysis. Future methods may choose to incorporate stricter filtering, and as our experiments have demonstrated, this not only helps in reducing data volume but also in improving the quality of search outputs. These issues may be alleviated by innovative computational strategies, such as taxonomic filters ^48^ or sweep deconvolution ^49^.

In light of technological development, the benefits of phylogenetic compression will grow over time. Currently, only a fraction of the world’s microbial diversity has been sequenced. However, as sequencing becomes more comprehensive, the tree of life will not change, thus enhancing the relative advantage of phylogenetic compression. We foresee its use ranging from mobile devices to large-scale distributed cloud environments and anticipate promising applications in global epidemiological surveillance ^50^ and rapid diagnostics ^51^. Overall, the phylogenetic compression of data structures has broad applications across computational biology and represents a fundamental design principle for future genomics infrastructure.

## METHODS

### Analysis of the decrease in bacteria BLAST searchability

#### Estimation of BLAST NT database size

The size of the BLAST NT database for the time period between 2002-01-01 and 2022-11-01 was estimated using five types of online resources, resulting in n=27 values. First, file sizes were manually recorded from the official NCBI website https://ftp.ncbi.nih.gov/blast/db/FASTA/ (n=11, between 2020-04-05 and 2022-11-01). Second, additional values were obtained from the snapshots of this website and its NCBI mirrors on http://web.archive.org (n=7, between 2012-10-11 and 2022-06-06). Third, archived versions of the NT database were found in diverse online repositories (n=3, between 2017-10-26 and 2021-01-15). Fourth, the NT database size was documented in a software documentation (n=1, 2013-12-03). Finally, the number of base pairs in the NT database was also documented in literature (n=5, between 2002-01-01 and 2010-01-01) (**Supplementary Table 8**). Conversion between the sizes of the GZip-compressed NT database and the corresponding total sequence lengths was performed using the 2.04 GZip bits per bp constant, estimated using the NT database as of 2022-06-20.

#### Estimation of NCBI Assembly database size

The number of bacteria in the NCBI Assembly database ^52^ (https://www.ncbi.nlm.nih.gov/assembly/) and their compressed size were estimated from the GenBank assembly summary file https://ftp.ncbi.nlm.nih.gov/genomes/genbank/bacteria/assembly_summary.txt (n=1,280,758 records, downloaded on 2022-11-02). The file was sorted according to the ‘`seq_rel_date`’ field and then used for calculating the number of published assemblies till a given date, aggregated per month. The total lengths of assemblies for the corresponding time points were estimated using the mean length of a bacterial genome assembly in the 661k collection (3.90 Mbp) and then converted to the estimated GZip size as previously. Although updates in the `assembly_summary.txt` file, such as the removal of old contaminated records, may influence the resulting statistics, a manual inspection during a several-months-long period showed only a minimal impact of these changes on the old statistics.

#### Comparison of BLAST NT and NCBI Assembly database sizes (**Fig 1a**)

To compare the sizes of two databases at the same time points, their respective functions were first interpolated in the logarithmic scale using piecewise linear functions from the data extracted above. The resulting interpolations were then used to calculate the estimated proportion of the sizes of NT and the bacteria in the NCBI Assembly database at regular intervals (monthly). Although minor inaccuracies might be present in the calculations (such as variations in the mean bacterial assembly or in the GZip-bits-per-bp conversion across different versions of the databases), these differences do not impact the overall exponential decrease of data searchability.

### Conceptual overview of phylogenetic compression

#### General overview

To organize input genomes into phylogenetic trees and compress/index them in a scalable manner, phylogenetic compression combines four conceptual steps.

#### Step 1: Clustering/batching (Fig. 1b(i))

The goal of this step is to partition genomes into batches of phylogenetically related genomes, of a limited size and diversity, that can be easily compressed and searched together using highly reduced computational resources. During downstream compression, indexing, and analyses, these individual batches are processed separately, and their maximum size and diversity can establish upper bounds on the maximum time and space necessary for processing a single batch. For instance, in the realm of *k*-mer aggregative methods (see an overview in ref ^15^), this corresponds to a matrix decomposition of a large *k*-mer annotation matrix into a series of small matrices that have both dimensions small, and analogically in the realm of dictionary compression, to reducing the input strings and dictionary sizes.

For microbes, clustering can be accomplished rapidly by metagenomic classification ^36^ applied to the raw reads or other methods for species identification. Microbial genomes in public repositories form distinct clusters, usually (but not always) corresponding to individual species ^35^, and metagenomic classification can assign individual genomes to these respective clusters, defined by the underlying reference database such as NCBI RefSeq ^36^. This requires only a constant time per dataset and can be fully parallelized, resulting thus in a constant-time clustering if sufficiently many computational nodes are available.

The obtained clusters are then reorganized into batches. First, too small clusters are merged, creating a special pseudo-cluster called dustbin, whose purpose is to collect divergent, weekly compressible genomes from sparsely sampled regions of the tree of life. Subsequently, the clusters that are too large – such as those corresponding to oversampled human pathogens (e.g., *S. enterica* or *E. coli*) – as well as the dustbin are then divided into smaller batches, to provide guarantees on the maximum required downstream computational resources per one batch. An additional discussion of batching is provided in **Supplementary Note 3**.

#### Step 2: Inference of a compressive phylogeny (Fig. 1b(ii))

In this step, the computed batches are equipped with a so-called *compressive phylogeny*, which is a phylogeny approximating the true underlying phylogenetic signal with sufficient resolution for compression purposes. If accurate inference methods such as RAxML ^53^ or FastTree 2 ^54^ cannot be applied due to the associated bioinformatics complexity or high resource requirements, phylogenies can be rapidly estimated via lighter approaches such as the Mashtree algorithm ^34^ (reimplemented more efficiently in Attotree, https://github.com/karel-brinda/attotree) instead, with only a negligible impact on the resulting compression performance (**Supplementary Fig. 5, Supplementary Note 1**).

#### Step 3: Data reduction/reordering (Fig. 1b(ii))

The compressive phylogenies obtained in the previous step serve as a template for phylogenetic reordering of individual batches. The specific form of reordering can vary depending on the specific data representations, intended applications, and method of subsequent compression or indexing. In principle, the reordering can occur in two directions: as a left-to-right genome reordering based on the topology of the compressive phylogeny, or as a bottom-up reduction of genomic content along the phylogeny (followed by left-to-right enumeration). Regardless of the specific form, this transformation is always reversible, thus sharing similarities with methods such as the Burrows-Wheeler transform ^21^.

#### Step 4: Compression or indexing using a calibrated low-level tool (**Fig. 1c**)

Finally, the reordered data are compressed or indexed using a low-level tool. At this stage, thanks to both phylogeny-based clustering and phylogeny-based reordering, the data are highly locally compressible, which enables to use of a wide range of general and specialized genome compressors/indexes. Nevertheless, it is crucial to ensure that the properties of the underlying algorithms and their parameters are closely tailored to the specific characteristics of the input data and their intended applications. For instance, to compress genomes in FASTA format, compressors based on Lempel-Ziv require the window/dictionary sizes to be large enough to span multiple genomes (**Supplementary Fig. 3a**), and general compressors also critically depend on FASTA being in a one-line format (**Supplementary Fig. 3b**). As a general rule, general compressors must always be carefully tested and calibrated for specific genomic data types, potentially requiring format cleaning and parameter calibration, whereas specialized genome compressors and indexers are usually pre-calibrated in their default setting and provided with well-tested configuration presets. While in many practical scenarios, individual batches are compressed/indexed separately, some protocols may involve merging reordered batches together to create a single comprehensive archive/index.

### The MiniPhy framework for phylogenetic compression

Here, we describe the specific design choices of our implementation of phylogenetic compression for assemblies and de Bruijn graphs. More information and relevant links, including specific tools such as MiniPhy and Phylign and the resulting databases, can be found on the associated website (https://brinda.eu/mof).

#### Clustering/batching

As genome collections encountered in practice can vary greatly in their properties as well as the available metadata, clustering is expected to be performed by the user. The recommended procedure is to identify species clusters using standard metagenomic approaches, such as those implemented in the Kraken software suite ^55^ (i.e., Kraken 2 ^56^ and Bracken ^57^ applied on the original read sets), as the obtained abundance profiles can also be used for quality control to filter out those samples that are likely contaminated. The next step is to divide the obtained genome clusters into smaller batches, analogically to the examples in **Supplementary Figure 1** and as discussed in more details in **Supplementary Note 3** (and the corresponding implementation in the MiniPhy package, see below). The order in which genomes are taken within individual clusters can impact the final compression performance; based on our experience, lexicographic order with accessions or ordering according to the number of distinct *k*-mers per genome provide surprisingly good performance as both of these approaches tend to group phylogenetically close genomes closer to each other. The protocol can be customized further to suit the performance characteristics of algorithms downstream, such as by adjusting the batch size or the parameters controlling the creation of dustbin batches (**Supplementary Note 3**). If the total size of a collection is small enough, the clustering/batching step may be skipped entirely and the entire collection treated as a single batch.

#### Inference of a compressive phylogeny

Users have the option to provide a custom tree generated by an accurate inference method such as RAxML ^53^. However, in most practical scenarios, such trees are not available, and MiniPhy then employs Attotree (https://github.com/karel-brinda/attotree), an efficient reimplementation of the Mashtree algorithm ^34^, to generate a compressive phylogeny through sketching. Both Mashtree and Attotree first use Mash ^58^ to estimate the evolutionary distances between all pairs of genomes, which are then used to infer a compressive phylogeny employing the Neighbor-Joining algorithm ^59,60^ as implemented in QuickTree ^61^. The distance computation in Mash is based on estimating the Jaccard indexes of the corresponding *k*-mer sets and then estimating the likely mutation rate under a simple evolutionary model ^62^. Finally, MiniPhy post-processes the obtained tree using standard tree-transformation procedures implemented in the ETE3 library ^63^, involving tree standardization, setting a midpoint outgroup, ladderization, and naming the internal nodes.

#### MiniPhy

(https://github.com/karel-brinda/miniphy). This is a central package for phylogenetic compression, including support for batching, and for calculating the associated statistics (see below). MiniPhy is implemented as a Snakemake ^64^ pipeline, offering three protocols for phylogenetic compression:

1. Compression of assemblies based on left-to-right reordering.
2. Compression of de Bruijn graphs represented by simplitigs ^33,65^ based on left-to-right reordering.
3. Compression of de Bruijn graphs through bottom-up *k*-mer propagation using ProPhyle ^28,29^.

In the third protocol, *k*-mer propagation is executed recursively in a bottom-up manner: at each internal node, the *k*-mer sets of the child nodes are loaded, their intersection computed, stored at the node, the intersection subtracted from the child nodes, and all three *k*-mer sets saved in the form of simplitigs ^33,65^; ProphAsm ^33^ performs all these operations. This process results in a progressive reduction of the *k*-mer content within the phylogeny in a lossless manner. Further details on this technique can be found in ref ^66^.

The output of each of the three protocols is a TAR file containing text files in their phylogenetic order, created from the corresponding list of files using the following command:

`tar cvf - -C $(dirname {input.list}) -T {input.list} --dereference`

For assemblies, these text files are the original assembly FASTA files, converted by SeqTK ^67^ to the single-line format with all nucleotides in uppercase (‘`seqtk` `seq` `-U {input.fa}`’). For simplitigs, the text files are EOL-delimited lists of simplitigs in the order as computed by ProphAsm, obtained from its output using the command ‘`seqtk` `seq` `{input.fa} | grep -v \>`’. The resulting TAR file is then compressed using XZ (‘`xz` `-9 -T1`’, see the section about calibration), and the resulting .tar.xz file distributed to users or further recompressed or indexed by other low-level tools, while preserving the underlying order.

#### MiniPhy statistics

For each of the three implemented protocols, MiniPhy generates a comprehensive set of statistics to quantify the compressibility of the batch, including: 1) *set* (the size of the *k*-mer set computed from all nodes of the compressive phylogeny), 2) *multiset* (the size of the *k*-mer multiset computed as a union of *k*-mer sets from individual nodes), 3) *sum_ns* (the total number of sequences), 4) *sum_cl* (the total sequence length), 5) *recs* (the number of records corresponding to individual nodes), and 6) *xz_size* (the size of the TAR file after XZ compression). The sizes of *k*-mer sets and multisets are determined from *k*-mer histograms computed by JellyFish 2 ^68^ (v2.2.10) using the commands:

`jellyfish count --threads {threads} --canonical --mer-len 31 --size 20M \`
`- -output {jf_file} {input}`

followed by

`jellyfish histo --threads {threads} --high 1000000 {jf_file}`

The computed statistics are used for calculating additional compression-related metrics, such as the number of bits per distinct *k*-mer or kilobytes per genome.

#### Phylogeny-explained redundancy

By comparing the sizes of *k*-mer sets and multisets before and after reduction by *k*-mer propagation along a compressive phylogeny, it is possible to quantify the proportion of the *k*-mer signal that is explained by the phylogeny. This yields the so-called *phylogeny-explained k-mer redundancy*, quantifying the proportion of redundant occurrences of canonical *k*-mers that can be eliminated through *k*-mer propagation, out of those potentially eliminable if the phylogeny perfectly explained the distribution of all the *k*-mers (i.e., every *k*-mer occurring only once after propagation and thus being associated with a single entire subtree):

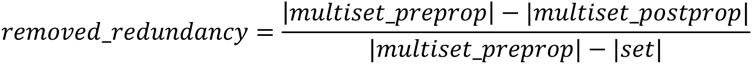

For collections comprising multiple batches, these variables refer to the global statistics, i.e., the sizes of set and multiset unions across all batches.

#### MiniPhy-COBS

MiniPhy-COBS (https://github.com/leoisl/miniphy-cobs) is a Snakemake ^64^ pipeline designed to create phylogenetically compressed ClaBS COBS indexes ^37^ (Classical Bit-sliced index) from assemblies already phylogenetically compressed by MiniPhy. ClaBS is a variant of COBS analogous to the original BIGSI data structure ^17^, using Bloom filters of the same size; this property is important for ensuring that the order of Bloom filters is preserved and that the neighboring Bloom filters are mutually compressible (**Supplementary Note 2**). The workflow for each batch involves three main steps:

1. Renaming input assemblies to align their lexicographic and phylogenetic orders within each batch,
2. Constructing COBS ClaBS indexes with:

`cobs classic-construct -T 8 {batch} {output}.cobs_classic`
3. Compressing the obtained indexes using:

`xz -9 -T1 -e --lzma2=preset=9,dict=1500MiB,nice=250`

#### Updated ProPhyle

To simplify the integration with MiniPhy for bottom-up *k*-mer propagation, a new version of ProPhyle ^28,29^ was released (v0.3.3.1, https://github.com/prophyle/prophyle). The main improvement compared to previous versions includes the possibility to stop after *k*-mer propagation, without proceeding to the construction of an FM-index, as such an index is unnecessary for phylogenetic compression using MiniPhy. The new version of ProPhyle is provided in the form of a Github release (https://github.com/iqbal-lab-org/cobs/releases) and pre-built packages on Bioconda ^69^.

### Overview of the five test microbial collections

#### GISP

The GISP collection comprises 1,102 draft assemblies of *N. gonorrhoeae* clinical isolates, collected in the US between 2000 and 2013 by the Centers for Disease Control and Prevention as part of the Gonococcal Isolate Surveillance Project (GISP) ^70^. These isolates had been sequenced using Illumina HiSeq and assembled using Velvet ^71^. The phylogenetic relationships among the isolates are known and had been determined using RAxML ^53^ after a correction for recombination by Gubbins ^72^. The GISP collection provides an example of a high-quality collection of draft genomes of a single low-diversity bacterial species, generated using a standardized sequencing and assembly protocol.

#### NCTC3k

The NCTC3k collection comprises 1,065 draft and complete assemblies of isolates of various bacterial species, derived from strains in the National Collection of Type Cultures (NCTC) collection and analyzed by Public Health England, the Wellcome Sanger Institute, and Pacific Biosciences as part of the NCTC 3000 Project ^73^ (https://www.culturecollections.org.uk/collections/nctc-3000-project.aspx). The isolates were sequenced using the PacBio Single Molecule, Real-Time (SMRT) DNA Sequencing technology, and assembled using automated pipelines. The assembled genomes are publicly available from the https://www.sanger.ac.uk/resources/downloads/bacteria/nctc/ website. The NCTC3k collection provides an example of a collection of high-quality, nearly complete genomes from diverse bacterial species.

#### SC2

The SC2 collection comprises 590,779 complete assemblies of SARS-CoV-2 isolates obtained from the GISAID database ^74^ as of 2021-05-18. These isolates were collected, sequenced, and assembled by various laboratories worldwide between 2020 and 2021 using various protocols. The phylogeny of the isolates is known and was computed by the sarscov2phylo software (https://github.com/roblanf/sarscov2phylo/, ref ^75^). The SC2 collection provides an example of a large collection of genomes of varying quality obtained from epidemiological surveillance of a single viral species at a global scale.

#### BIGSIdata

The BIGSIdata collection comprises 425,160 cleaned de Bruijn graphs representing nearly all bacterial and viral isolates available in the European Nucleotide Archive (ENA) as of December 2016 ^17^. These isolates had originally been collected and sequenced by various laboratories worldwide, deposited as raw-read data or genome assemblies to repositories synchronized with the ENA (ENA, NCBI SRA, and DDBJ Sequence Read Archive), and later downloaded and transformed into cleaned de Bruijn graphs using McCortex ^40,76^ (k=31) by the European Bioinformatics Institute (EBI). The resulting graphs were provided on an HTTP/FTP website (http://ftp.ebi.ac.uk/pub/software/bigsi/nat_biotech_2018), along with metadata on Figshare ^77^, although not all of the original 447,833 graphs could be retrieved in this study (see below). The BIGSIdata collection provides an example of a large and diverse collection of bacterial and virus isolates, collected and sequenced across the globe using various sequencing technologies and all provided in a unified graph representation.

#### 661k

The 661k collection comprises 661,405 draft assemblies of all Illumina-sequenced bacterial isolates present in the ENA as of November 2018 ^16^. These isolates had originally been collected and sequenced by various laboratories worldwide, and their raw-read data deposited to repositories synchronized with the ENA (ENA, NCBI SRA, and DDBJ Sequence Read Archive). The assemblies were generated using a single unified pipeline (https://github.com/iqbal-lab-org/assemble-all-ena) based on Shovill (https://github.com/tseemann/shovill) by EBI, and provided on an HTTP/FTP website (https://ftp.ebi.ac.uk/pub/databases/ENA2018-bacteria-661k/), along with metadata on FigShare ^78^.

The 661k collection provides an example of a large and diverse collection of assembled bacterial isolates, collected and sequenced across the globe using a single sequencing technology, i.e., the state-of-the-art of the short read-assembly era.

Basic characteristics of the five test collections, including the original file size, number of samples, species count, and the number of distinct *k*-mers, are provided in **Supplementary Table 1**.

### Acquisition of the test collections

#### GISP

The GISP collection was obtained from the https://github.com/c2-d2/rase-db-ngonorrhoeae-gisp repository (version 04a132c) as published in ref ^51^. The assemblies (n=1,102) were obtained from the “isolates/contigs” subdirectory of Github repository (containing the original genomes including the plasmids), and the associated RAxML phylogenetic tree was downloaded from the “tree/” subdirectory of the same repository. The original data had originally been analyzed in ref ^70^ and provided for download on Zenodo ^79^.

#### NCTC3k

The assemblies were obtained in the GFF format from ftp://ftp.sanger.ac.uk/pub/project/pathogens/NCTC3000 by

`wget -m -np -nH --cut-dirs 3 –retr-symlinks \`
ftp://ftp.sanger.ac.uk/pub/project/pathogens/NCTC3000.

The obtained files were converted them to the FASTA format by any2fasta (https://github.com/tseemann/any2fasta, v0.4.2) parallelized by GNU Parallel ^80^ and uploaded to Zenodo (ref ^81^, http://doi.org/10.5281/zenodo.4838517). The number of species in the collection was determined based on the data provided in the main Sanger/Public Health England assembly table for NCTC 3000 (https://www.sanger.ac.uk/resources/downloads/bacteria/nctc/, retrieved on 2022-09-14). The HTML table was manually exported to XLSX and used to construct a translation table from NCTC accession numbers to corresponding species. The accessions of the assemblies in our collection were then extracted from file names, translated to species, and the species counted. Overall, this resulted in n=1,065 assemblies of 259 species.

#### SC2

The SARS-Cov-2 data were downloaded from the GISAID website (https://www.gisaid.org/, as of 2021-05-18) in the form of an assembly file (‘`sequences_fasta_2021_05_18.tar.xz`’, n=1,593,858) and a Sarscov2phylo phylogeny ^82^ (‘`gisaid-hcov-19-phylogeny-2021-05-11.zip`’, n=590,952). After converting both files to the same set of identifiers and removing isolates with missing data, we obtained n=590,779 genome assemblies organized in a phylogenetic tree.

#### BIGSIdata

The BIGSI collection data ^17^ were downloaded from the associated FTP (http://ftp.ebi.ac.uk/pub/software/bigsi/nat_biotech_2018/), including cleaned de Bruijn graphs, taxonomic information inferred using Kraken ^26^, and abundance reports computed using Bracken ^57^. The download was done using RSync in groups corresponding to individual EBI prefixes (e.g., DRR000) by

`rsync -avP --min-size=1 --exclude ’*stats*’ --exclude ’*uncleaned*’ \`
`- -exclude ’*bloom*’ --exclude ’*log*’ \`
`rsync://ftp.ebi.ac.uk/pub/software/bigsi/nat_biotech_2018/ctx/{prefix}`

The prefixes were organized into 15 groups of at most 100 prefixes each, and the groups were processed individually in succession on a research computing cluster, with a parallelization using Slurm and jobs deployed using Snakemake ^64^ (between 2020-08-01 and 2020-09-15). From the downloaded McCortex files, unitigs were extracted using McCortex:

`bzcat -f {input} | mccortex31 unitigs -m 3G –`

Only those graphs with an uncorrupted McCortex file, Bracken information available, unitigs of total length ≥2 kbp with ≤15 M distinct *k*-mers, and with no file system error encountered were used in the subsequent processing. This resulted in n=425,161 de Bruijn graphs (out of the original n=463,331 genomes from the FTP or n=447,833 genomes reported in ref ^17^).

#### 661k

The 661k collection was downloaded in March 2021 from the official FTP repository specified in ref ^16^, using RSync by

`rsync-avp rsync://ftp.ebi.ac.uk/pub/databases/ENA2018-bacteria-661k/Assemblies/{pref}`

The command was run for individual prefixes ranging from 000 to 661, which resulted in n=661,405 .fa.gz files.

### Calibration and evaluation of phylogenetic compression

#### Calibration of XZ as a low-level tool for phylogenetic compression (Supplementary Fig. 3)

The compression performance of GZip, BZip2, and XZ was evaluated using the GISP collection, converted to the single-line FASTA format and with genomes sorted left-to-right according to the Mashtree phylogeny. For each compressor, the compression was performed with a range of presets and always with a single thread. To evaluate the compression performance with large resources available, two additional manually tuned modes with larger dictionaries, denoted by ‘M’ and ‘MM’, were added to the XZ benchmark, corresponding to the parameters

`- -lzma2=preset=9,dict=512MiB`

and

`- -lzma2=preset=9,dict=1500MiB,nice=250`

respectively.

To evaluate the impact of different line lengths on the compression, the source FASTA was reformatted for different lengths using SeqTK ^67^ and compressed using XZ by

`seqtk seq -l {line_length} | xz -9 -T1`

#### Comparison of scaling modes (Supplementary Fig. 4)

The SC2 collection was provided in the left-to-right order according to Sarscov2phylo phylogeny. The genomes were progressively uniformly subsampled, stored as EOL-separated lists of sequences (without sequence headers), and then compressed using individual compressors, namely: 1) XZ: ‘`xz` `-9 -T1`’, 2) BZip2: ‘`bzip2` `--best`’, 3) GZip: ‘`gzip` `-9`’, and 4) Re-Pair ^83,84^ (https://github.com/rwanwork/Re-Pair, version as of 2021-10-26):

`repair -v -I {inp_seqs}; tar cf {inp_seqs}.tar {inp_seqs}.prel {inp_seqs}.seq`

As Re-Pair did not provide sufficient scalability for the entire SC2 data set and the implementation suffered from various bugs, the Re-Pair sub-experiment was limited only to n≤70k, the integrity of the output files always verified via their decompression and line counting, and all archives lacking integrity were discarded from the subsequent analysis.

The scalability comparisons for the NCTC3k and GISP collections were performed analogically, but using MiniPhy (commit ‘`41976c7`’) and with sequence headers preserved. The order of all assemblies was first randomized by ‘`sort` `-R`’ and the individual sub-samplings for compression then generated as prefixes of this randomized list. The size comparisons were made based on the .tar.xz output file of the pipeline, as well as additional files obtained via their recompression by GZip and BZip2 with the same parameters as above.

#### Order comparison (Supplementary Fig. 5)

The SC2 collection was put into three different orderings: the original ordering (corresponding to the lexicographical ordering by sequence names), the left-to-right ordering of the phylogeny, and a randomized order. In all cases, a custom Python script using BioPython ^85^ was used to order the FASTA file and remove sequence names, and its output was compressed by the XZ compressor using 1 thread and the best preset (‘`xz` `-T1 -9`’). The comparisons for GISP and NCTC3k was performed analogically, but with sequence headers preserved.

#### Summary of MiniPhy calibration

XZ with the parameters ‘`xz` `-9 -T1`’ was chosen as the default compression procedure for MiniPhy, and Mashtree ^34^ as the default method for inferring compressive phylogenies. These choices were done based on the observations that the most popular method, GZip, always performed poorly for bacteria, although provided a moderate compression performance for viruses. On the other hand, XZ achieved steep compression curves for low-diversity collections, with compression ratio improving by one order per one order increase of the number of genomes, for both viruses and bacteria. NCTC3k as a high-diversity collection was weakly compressible even with the best approaches (<1 order of magnitude of compression after a 3 orders-of-magnitude increase of the number of genomes). One of the best available (but still highly experimental) grammar-based compressors, Re-Pair ^83,84^, achieved a similar asymptotic behavior as XZ, indicative of the potential of grammar compressors for phylogenetic compression to provide random access, but its usability remains experimental. Phylogenetic reordering boosted compression substantially for both low-and high-diversity collections (reduction in size between 38% and 67% compared to random orders). Finally, compressive phylogenies computed using Mashtree ^34^ provided nearly equal compression performance as an accurate approach using RAxML ^53^.

### Phylogenetic compression of the BIGSIdata collection of de Bruijn graphs

#### Clustering and batching

For every sample, the output of Kraken ^26^ and Bracken ^57^ were extracted from the downloaded data as provided in the online FTP repository (http://ftp.ebi.ac.uk/pub/software/bigsi/nat_biotech_2018/ctx/) in the Bracken files (‘`{accession}.ctx_braken.report`’) as the previously identified most prevalent species (corresponding to the row with the highest value of the ‘`fraction_total_reads`’ column). Clustering and batching then proceeded as depicted in **Supplementary Fig. 1** and further commented in **Supplementary Note 3**, with genomes being sorted according to the number of *k*-mers before their partitioning into batches. Overall, the genomes of the 1,443 identified species (clusters) were partitioned into 568 regular batches and 6 dustbin batches, resulting in a total of 574 batches.

#### Phylogenetic compression

Phylogenetic compression was performed twice, with slightly different workflows.

First, phylogenetic compression proceeded manually, via a workflow whose modified version was later implemented in MiniPhy. For individual batches, compressive phylogenies were computed using Mashtree ^34^ with the default parameters. The resulting trees and McCortex unitig files were then used as input for ProPhyle (v0.3.3.0) to propagate *k*-mers along the phylogenies, compute simplitigs ^33^, and merge the output FASTA files into a single one by

`prophyle index -k 31 -A -g {dir_genomes} {tree} {batch_name}`

The resulting FASTA files produced by ProPhyle (called ‘index.fa’) were converted into the single-line format using SeqTK ^67^ and compressed using XZ by

`seqtk seq {prophyle_index_fa} | xz -9 -T8`

The resulting files occupied 74.4 GB and were deposited on https://doi.org/10.5281/zenodo.4086456 and https://doi.org/10.5281/zenodo.4087330. Support for this version of the data set was incorporated into De-MiniPhy-BIGSIdata (see below), and the correctness of the end-to-end protocol and of the resulting files was validated by De-MiniPhy-BIGSIdata and subsequent *k*-mer counting using kc-c3 (https://github.com/lh3/kmer-cnt, commit ‘`e257471’`). The obtained *k*-mer counts were compared to those obtained from the original McCortex files (from the total length and count of unitigs); all *k*-mer counts were equal with the exception of 4 samples with 17–26 more reported *k*-mers after decompression.

Second, an analogical version of the propagated simplitig files, but without sequence headers and with compression using a single thread only, was later created using the MiniPhy pipeline and resulted in files occupying in total 52.3 GB that were subsequently deposited on https://doi.org/10.5281/zenodo.5555253.

#### Decompression of BIGSIdata de Bruijn graphs

To decompress de Bruijn graphs from the files obtained by *k*-mer propagation, all *k*-mers along all root-to-leaf paths need to be collected. We implemented this specifically for BIGSIdata in a Python package called De-MiniPhy-BIGSIdata (https://github.com/karel-brinda/De-MiniPhy-BIGSIdata). The program downloads individual data files from Zenodo from the accessions above (the first version of the dataset) and reconstructs the original *k*-mer sets using the following procedure. First, it decompresses the XZ file of a given batch, splits it according to files corresponding to individual nodes of the compressive phylogeny, recompresses individual nodes using GZip parallelized by GNU Parallel ^80^, and for all leaves (genomes) it reconstructs the corresponding *k*-mer sets by merging all GZip files along the corresponding root-to-leaf paths using the Unix cat command. From the obtained output FASTA files, de Bruijn graphs can be easily reconstructed by standard tools such as BCALM2 ^86^.

#### Comparison to the original compression protocol

As the samples in our BIGSIdata collection do not fully correspond to the data that were used in the original publication of BIGSI ^17^, we recalculated the size statistics of the published McCortex files of our graphs based on the FTP list-off files as provided within individual subdirectories of http://ftp.ebi.ac.uk/pub/software/bigsi/nat_biotech_2018/ (as of 2021-08-27). These were downloaded per individual prefix directories recursively using wget by

`wget -nv -e robots=off -np -r -A .html \`
http://ftp.ebi.ac.uk/pub/software/bigsi/nat_biotech_2018/ctx/{prefix}/

The corresponding parallelized Snakemake pipeline was run on a desktop computer. This resulted in a table containing 484,463 files, out of which 162,645 were BZip2-compressed. The individual file records were compared with the list of accessions of files that were previously retrieved and sorted in our BIGSIdata collection, and the volume of the source graphs on FTP calculated to be 16.7 TB.

#### Comparison to Metagraph ^38^

The size of the phylogenetically compressed BIGSIdata collection was compared to the size of an analogous Metagraph index from the original paper ^38^, based on the statistics in Table 1 and Supplementary Table 1 therein (the SRA-Microbe collection): n=446,506 indexed datasets, 39.5 G canonical *k*-mers (with the same *k*-mer size *k*=31), and the size of the annotated de Bruijn graph being 291 GB (graph 30 GB + annotations 261 GB). This index was constructed from the same datasets as in the original BIGSI paper ^17^ but using a slightly different computational methodology. Consequently, the index of Metagraph contained approximately 4% fewer distinct canonical *k*-mers (k=31) compared to BIGSIdata as used in this paper. To compare the two compression approaches (MiniPhy with bottom-up *k*-mer propagation and XZ as a low-level tool vs. Metagraph), both applied to the similar but different input data, we used the number of bits per distinct *k*-mer as the statistic for comparison, which was found to be 10.2 and 58.9, respectively. Therefore, the MiniPhy compression was more efficient by an estimated factor of 5.78. We note that phylogenetic compression could be directly embedded into Metagraph (by imposing the phylogenetic order of columns during index construction), which may help to further reduce its index size.

### Phylogenetic compression of the 661k assembly collection

#### Clustering and batching

Species clusters were identified based on the most prevalent species in the sample as identified using Kraken 2 ^56^ and Bracken ^57^ from the original raw-read data; i.e., based on the ‘`V2`’ column in the ‘`File1_full_krakenbracken.txt`’ file of the supplementary materials of ref ^16^. The creation of the dustbin pseudo-cluster and formation of individual batches proceeded by the steps documented in **Supplementary Fig. 1** and as later implemented directly within MiniPhy, with genomes pre-sorted lexicographically according to ENA accessions.

#### Phylogenetic compression using MiniPhy

The obtained batches were compressed using the MiniPhy pipeline as described above; i.e., compressive phylogenies were computed using Mashtree ^34^ and used for 1) left-to-right reordering of the assemblies, 2) left-to-right reordering of simplitigs of the corresponding de Bruijn graphs, and 3) bottom-up *k*-mer propagation and simplitig computation by ProPhyle; while in all cases storing the simplitigs and assemblies as text and FASTA file, respectively, followed by a compression by ‘`xz` `-9 -T1`’. The compressed assemblies were deposited on https://doi.org/10.5281/zenodo.4602622.

#### Calculations of the statistics

All the statistics used in the plots and tables were calculated based on the numbers obtained from MiniPhy. Additionally, the total number of *k*-mers was calculated using JellyFish ^68^ (v2.2.10) by

`jellyfish count --mer-len 31 --size 200G --threads 32 \`
`- -output kmer_counting.jf --out-counter-len=1 --canonical`

which resulted in 44,349,827,744 distinct *k*-mers (28,706,296,898 unique *k*-mers) for the 661k collection and in 35,524,194,027 distinct *k*-mers (22,904,412,202 unique *k*-mers) for the 661k-HQ collection (as described below). The files uploaded to https://doi.org/10.5281/zenodo.4602622 are higher by approximately 0.2 GB (approx. 0.7% of the total size) compared to the value **Supplementary Table 3** as the Zenodo submission was done with an older version of compressive phylogenies without their post-processing.

#### Recompression using MBGC

Individual phylogenetically compressed batches from the previous step were converted to single FASTA files by ‘tar -xOvf {input.xz}*’* and then compressed using MBGC ^18^ (v1.2.1) with 8 threads and the maximum compression level by

`mbgc -i {input.fa} -c 3 -t 8 {output.mbgc}`

#### Compression in the lexicographic order

Data in ENA and other similar repositories have identifiers assigned in the order in which they are uploaded, individual uploads typically proceed by uploading entire projects, and these typically involve phylogenetically very close genomes. For instance, genomes from a study investigating a hospital outbreak often occupy a range of accessions. Therefore, lexicographically sorted genomes from ENA may be used as an approximation of phylogenetic compression. To compare the compressibility of the 661k collection in the ENA accession lexicographic order to the full phylogenetic compression, we streamed the genomes from the main collection file provided on http://ftp.ebi.ac.uk/pub/databases/ENA2018-bacteria-661k/661_assemblies.tar, decompressed them on-the-fly, converted them to the one-line FASTA format using SeqTK ^67^, and compressed them using XZ with 32 threads by

`pv 661_assemblies.tar | tar -xOf - | gunzip -c | seqtk seq | xz -9 -T32`

### Phylogenetic compression of the 661k/661k-HQ k-mer indexes

#### The 661k-HQ collection

To reduce biases in *k*-mer matching, a high-quality variant of the 661k collection, called 661k-HQ, was constructed from the 661k collection by excluding genomes that had not passed quality control in the original study ^16^ (3.7% of the genomes). For simplicity, the batches and genome orders in 661k-HQ were kept the same as in 661k.

#### Phylogenetic compression of the 661k/661k-HQ COBS indexes

COBS indexes for the 661k and 661k-HQ collection were constructed per batch using the MiniPhy-COBS pipeline (see the MiniPhy-COBS section), which produces the ClaBS variant of the index with all Bloom filters of the same size sorted left-to-right according to the phylogeny, and compresses them using XZ.

#### Comparisons to the compact COBS indexes

The compact variant of the COBS index (default in COBS), based on adaptive adjustments of Bloom filter sizes through subindexes of different heights, was used as a baseline in our comparisons. For the 661k collection, we used the original index as provided (http://ftp.ebi.ac.uk/pub/databases/ENA2018-bacteria-661k/661k.cobs_compact, retrieved on 2022-09-08, 937 GB). For building a COBS index for 661k-HQ, we used the same construction protocol as in ref ^16^. Both indexes were then compressed on a highly performant server by XZ using 32 cores (‘`xz` `-9 -T32`’).

All of the obtained data points are provided in **Supplementary Table 5**.

### Phylign pipeline for alignment against all pre-2019 bacteria from ENA

#### Overview

The Phylign pipeline (https://github.com/karel-brinda/phylign) uses phylogenetically compressed assemblies (661k) and COBS indexes (661k-HQ) as described above to align queries against the entire 661k-HQ collection in a fashion similar to BLAST (**Supplementary Note 4**). The search procedure consists of two phases: matching the queries against the *k*-mer indexes using COBS ^37^ to identify the database’s most similar genomes for each query, followed by an alignment of the queries to their best-matching genomes using Minimap 2 ^41^. Phylign is developed as a Snakemake ^64^ pipeline, using Bioconda ^69^ for an automatic software management and the standard Snakemake resource management ^64^ to control the CPU cores assignments and limit RAM usage according to user-specified parameters. Upon its first execution, Phylign downloads its phylogenetically compressed reference database from the Internet (102 GB), consisting of 29.2 GB of assemblies and 72.8 GB of COBS indexes.

#### Matching

The matching step involves *k*-mer matching of all user queries against the entire 661k-HQ database using a modified version of COBS (v0.3, see below), based on the principle that the number of *k*-mer matches between a genome and a query correlates with the alignment score ^87^. Each phylogenetically compressed COBS index is decompressed in memory and queried for the input user sequences, reporting all matches between the queries and genomes in the current batch with a sufficient (user-specified) proportion of matching *k*-mers. The computed matches are then aggregated across all batches, and for each query, only a (user-specified) number of best matches, plus ties, are retained and passed to the subsequent alignment step. Matching is parallelized by Snakemake, with the number of threads for each COBS instance adjusted based on batch size.

#### Alignment

For each batch independently and fully in parallel, Phylign then iterates over the phylogenetically compressed genome assemblies, and if a given genome has at least one match passed from the matching phase, it builds on-the-fly, in memory, a new Minimap 2 ^41^ (v2.24) instance for this genome and aligns all relevant queries to this genome, while saving Minimap 2 outputs in a batch-specific output file. Once all batches are processed, the resulting alignments are aggregated and provided to the user in a modified SAM format ^88^.

#### Performance characteristics

The total matching time is primarily driven by the time complexity of COBS, with decompression accounting for less than 2 CPU hours (**Supplementary Fig. 9**). In the alignment step, decompression requires less than 1.5 CPU hours (**Supplementary Fig. 9**), and the remainder of the time is primarily driven by the time to create a new Minimap2 instance (estimated 0.3 CPU seconds per instance in the current implementation). If the queries are long and Minimap 2 is used with a sensitive preset, the actual Minimap 2 alignment time becomes the main time component (e.g., in the plasmid experiment in **Supplementary Tab. 6**).

#### Updated COBS

To integrate COBS into Phylign, new versions of COBS ^37^ were created (v0.2, v0.3, https://github.com/iqbal-lab-org/cobs). The updates include support for macOS, streaming of indexes into memory, and multiple bug fixes. The new versions of COBS are provided in the form of Github releases (https://github.com/iqbal-lab-org/cobs/releases) and pre-built packages on Bioconda ^69^.

#### Benchmarking of the decompression time

Decompression times were evaluated on the same desktop computer as the alignment experiments, separately for the phylogenetically compressed assemblies vs. COBS indexes and for in-memory decompression (‘`xzcat` `{file}` > `/dev/null`’) vs. on-disk decompression (‘`xzcat` `{file}` > `{tmpfile}`’), resulting in four experiments. Within each experiment, decompression was parallelized using GNU Parallel (‘`parallel` `-L1 -v –progress`’), with time measured using GNU time both for the whole experiment and for each batch in a given compressed representation.

### Evaluating Phylign

#### Overview of the benchmarking procedure

The search using Phylign was evaluated on three datasets, representative of different query scenarios: a database of antibiotic resistance genes, a database of plasmids, and an Oxford nanopore sequencing experiment. In all cases, the search parameters – including the number of hits of interest, the COBS *k*-mer threshold, and the Minimap preset – were tailored to each specific query type. The experiments were conducted on an iMac with a Quad-Core Intel CPU i7, 4.2 GHz with 4 physical (8 logical) cores and 42.9 GB (40 GiB) RAM.

#### Time measurements

The wall clock and CPU time were measured using GNU time and calculated as real and usr+sys, respectively. The measurements were done for the matching and alignment steps separately.

#### Memory measurements

We have not found any reliable way of measuring peak memory consumption on macOS: both GNU time and the psutil Python library were significantly underestimating the memory footprint of our Snakemake pipeline. Therefore, we performed additional measurements on a Linux cluster using the SLURM job manager, using jobs allocated with a configuration similar to the parameters of our iMac computer. For ‘`max_ram_gb`’ set to 30 GB, we observed a peak memory consumption of 26.2 GB, thus by 12.7% lower compared to the specified maximum. Such a discrepancy is expected because the ‘`max_ram_gb`’ parameter defines an upper bound for the Snakemake resource management ^64^, representing the worst-case scenario for parallel job combinations.

#### Resistance genes – ARGannot

The resistance genes search was performed using the ARG-ANNOT database ^42^ comprising 1,856 genes/alleles, as distributed within the SRST2 software toolkit ^89^ (https://github.com/katholt/srst2/blob/master/data/ARGannot_r3.fasta, retrieved on 2022-07-24). The search parameters were set to require a minimum of 50% matching *k*-mers, with 1,000 best hits plus ties taken for every gene/allele query. Alignment was performed with the Minimap preset for short reads (‘`sr`’).

#### Plasmids – the EBI plasmid database

The list of EBI plasmid was downloaded from the associated EBI website (https://www.ebi.ac.uk/genomes/plasmid.details.txt, retrieved on 2022-04-03), and individual plasmids were subsequently downloaded from the ENA using curl and GNU parallel ^80^. The search parameters were set to require at least 40% matching *k*-mers (the threshold previously used in ref ^17^), with 1,000 best hits plus ties taken for every plasmid. Alignment was performed with the Minimap preset for long, highly divergent sequences (‘`asm20`’).

#### Oxford Nanopore reads

The ERR9030361 experiment, comprising 159k nanopore reads from an isolate of *M. tuberculosis*, was downloaded from SRA NCBI. The search parameters were set to require at least 40% matching *k*-mers, with 10 best hits plus ties taken for every read. Alignment was performed with the Minimap preset for nanopore reads (‘`map-ont`’).

#### Comparison to BIGSI

As we were unable to reproduce the original plasmid search experiment ^17^ with BIGSI on our iMac computer (due to the required database transfer of 1.43 TB over an unstable FTP connection), we used the values provided in the original publication ^17^. To ensure a fair comparison, we focused on evaluating the total CPU time (sys+usr) and verified that our parallelization efficiency was close to the maximal one (680% out of 800% possible achieved, based on the values in **Supplementary Table 7**).

## DATA AVAILABILITY STATEMENT

All data supporting the findings of this study and the developed software are available within the paper and its Supplementary Materials (**Supplementary Table 4**).

## ACKNOWLEDGEMENTS

This work was partially supported by the NIGMS of the National Institutes of Health (R35GM133700), the David and Lucile Packard Foundation, the Pew Charitable Trusts, and the Alfred P. Sloan Foundation. R.C. was partially supported by the European Union’s Horizon 2020 research and innovation programme (grants agreements No. 872539, 956229, and 101047160) and ANR Transipedia, SeqDigger, Inception, and PRAIRIE grants (ANR-18-CE45-0020, ANR-19-CE45-0008, PIA/ANR16-CONV-0005, ANR-19-P3IA-0001). Portions of this research were conducted on the O2 high-performance compute cluster, supported by the Research Computing Group at Harvard Medical School, and on the GenOuest bioinformatics core facility (https://www.genouest.org).

## Supplementary notes

### Supplementary Note 1. Stability of phylogenetic compression

The overall performance of phylogenetic compression stems from a combination of trade-offs between the individual layers of a given phylogenetic compression protocol (such as for assemblies, de Bruijn graphs, or *k*-mer indexes). These layers include the specific clustering and batching strategy, compressive phylogeny inference, and the low-level compression/indexing technique.

#### Clustering

Clustering can be performed using various direct or indirect methods. All these methods expected to identify similar clusters thanks to the pronounced species structure across public microbial isolate dataset ^35^. However, both classes of approaches have specific caveats that may downgrade the resulting compression performance.

##### Caveats of indirect approaches

When clustering is based on species identification by Kraken or other LCA-based classifiers, clustering might be impacted by the loss of resolution due to reference database growth ^90^. While this is unlikely to significantly affect phylogenetic compression performance with collections akin to 661k (where phylogenetically related genomes would still be clustered together, although under biological incorrect species names); a carefully analysis of the data structure will be necessary for atypical collections, such as those comprised of metagenome-assembled genomes.

##### Caveats of direct approaches

Direct clustering methods, now feasible at the scale of millions genomes ^91^, are contamination-oblivious and may thus be sensitive to various contamination patters (see, e.g.,, the discussion of *C. difficile* in ref ^91^). Contamination is very common in public genomic datasets, and if not properly controlled by metagenomic profiling or other quality control techniques, it can impede both downstream compression and search.

#### Batching

For 661k and BIGSI data, batching has been implemented heuristically, with lexicographic preordering based on accessions, to ensure that genomes sequenced around the same time would, within the same species cluster, be batches together. An alternative pre-sorting strategy, based on the number of *k*-mers in a given dataset, was tested for BIGSIdata (data not shown), and led to mostly comparable results.

#### Compressive phylogeny

In most scenarios, compressive phylogeny is used for within-batch reordering of either assemblies directly or of columns corresponding to individual genomes in case of *k*-mer indexes. When clustering and batching are done correctly and a robust low-level compressor used (e.g., XZ), such reordering by itself provides a moderate improvement (30–55% reduction, see **Supplementary Fig. 5**). Nevertheless, the impact is much stronger with less advanced compression techniques; for instance, run-length encoding (RLE) applied to *k*-mer matrices improves by up to an order of magnitude when the columns are reordered according to phylogenies (data not shown). When considering different approaches to compute phylogenies, even sketching combined with neighbor joining provides a sufficient resolution; Mashtree yields nearly as good compression results as full-scale methods for phylogenetic inference, such as RAxML ^53^. Differences in the resulting compression ratios are relatively minor, with RAxML phylogenies showing a slight advantage in Lempel-Ziv-based compression on assemblies over Mash trees (**Supplementary Table 3**), and conversely, Mash trees exhibit slightly better performance in compressing de Bruijn graphs or *k*-mer sets (**Supplementary Table 2, 3**).

#### Low-level compressor or indexer

The final performance of phylogenetic compression is significantly influenced by the capabilities of the used low-level compressor or indexer. For dictionary compressors, an essential parameter is the dictionary size or the window size (**Supplementary Fig. 3a, Supplementary Fig. 4**), which disqualifies many popular compressors, including gzip and bzip2. For general compressors applied to assemblies, a crucial factor is converting FASTA to the one-line format (**Supplementary Fig. 3b**). There are also notable differences in compression speed: compressing a single batch of assemblies using XZ might require up to several hours (albeit with rapid decompression), while MBGC (v2.0) requires approximately ten minutes per batch.

### Supplementary Note 2. Genome-similarity-preserving representations in phylogenetic compression

As a prerequisite for phylogenetic compression, it is fundamental to assume that the core genome representations preserve similarity. Informally, this means that little changes in the input genome lead to only little changes in its representation, ensuring that closely phylogenetically related genomes have highly mutually compressible representations. Although the similarity-preserving property can be rigorously defined in specific cases using mathematical formalism including specific input and output distances and embeddings, we adopt a more conceptual perspective to maintain a broader view.

#### Examples of genome-similarity-preserving representations

- **Complete genomes assemblies.** Complete genome assemblies precisely reflect the sequence of nucleotides in DNA molecules, with single mutation events resulting in single changes in the assembly.
- **Draft genome assemblies.** Similar to complete assemblies, but may not fully resolve repetitive regions, leading to a fragmented assembly. In contrast to complete assemblies, a single evolutionary event might induce a more substantial change in the representation. For example, a mutation in a previously non-resolvable repetitive region could make it resolvable by turning an exact repeat into an inexact one. Nevertheless, such events are rather infrequent, and for many compression techniques (e.g., those based on Lempel-Ziv), the distance between the two representations remains minimal.
- **Burrows-Wheeler Transform of assemblies.** The Burrows-Wheeler transform ^21^ is characterized by its locality, in the sense that a localized change in the input induces only a localized change in the BWT-transformed string ^92^.
- ***k*-mer spectrum.** Changing, deleting, or inserting one nucleotide in the genome alters the *k*-mer spectrum by the removal and addition of up to 2*k*+2 *k*-mers.
- **MinHash sketches.** The addition or removal of a *k*-mer to or from a spectrum may lead to the replacement of one hash value by a smaller or larger one, respectively, and such a replacement happens only with a very low probability. Therefore, sketches of similar genomes are either identical or very similar.
- **Minimizer de Bruijn graphs.** These combine properties of de Bruijn graphs and minimizers, with the genome-similarity-preserving property following naturally from this combination.
- **Bloom filters of fixed size.** The addition or removal of element to or from a set always alters the Bloom filter by a maximum of *m* bits, where *m* is the number of hash functions; therefore, a small change in the genome results only in a small change in the corresponding fixed-size Bloom filter.

#### Examples of representations that are not similarity-preserving

- **Bloom filters with adaptive sizes.** Adaptive size adjustment (such as implemented in COBS’s default strategy ^37^, which uses smaller Bloom filters for smaller genomes), disrupts similarity preservation. For instance, an event such as an acquisition of a plasmid by an *E. coli* strain may cause the Bloom filter to expand, reflecting an increase in genome and *k*-mer set size, altering also the underlying hash functions (or the associated modulo function). In consequence, adaptive-size Bloom filters of even closely related genomes can be very dissimilar. As a result, we did not use the COBS default strategy, but forced it to use the same size of Bloom filters for all genomes in a given batch.

### Supplementary Note 4. Core principles of the MiniPhy batching approach

The batching approach used in this paper, as summarized for the 661k and BIGSIdata collections in **Supplementary Fig. 1**, is based on the following principles. At its core, phylogenetic compression involves the phylogenetic reordering of input data. For large collections, this process entails partitioning genomes into batches that adhere to specific constrains on certain characteristics, and then reordering them phylogenetically based on compressive phylogenies.

To ensure the essential guarantees from the paper, and to maximize the batches’ usability across combinations of tools and in diverse application use cases, the batches are required to have following properties:

1. **An upper-bounded compressed size** – to guarantee easy internet transmission, even over unreliable networks.
2. **A lower-bounded compressed size** – to limit the negative effects of excessively unbalanced batches in workflow managers such as Snakemake and Nextflow and in resource allocation systems such as Slurm.
3. **An upper-bounded decompressed size** – to minimize the maximum memory required per batch in downstream data analysis and to facilitate the parallel processing of multiple batches in memory-constraint environments.
4. **An upper-bounded number of genomes per batch** – to establish a limit on the time required per batch for phylogenetic inference and for downstream data analyses.
5. **Optimization for maximal compression ratio within these constraints** – to minimize the overall necessary data transmission over the Internet and within a computer (e.g., from disk to RAM).

On a mathematical level, these constrains lead to interesting optimization problems that may be formalized and solved by techniques such as integer linear programming or answer set programming, in combinations with techniques for estimating data compressibility via measures such as the size of minimal string attractors ^93^, factor complexity ^94^, or the δ measure ^95^.

However, for simplicity, our approach used in MiniPhy is empirical, informed by the following observations about bacterial genomes and the structure of ENA:

1. **Constrained genome size range.** For bacteria, their genome size can be assumed to fit within a range of one order of magnitude, typically 1 Mbp to 10 Mbp (see the principles behind BIGSI ^17^).
2. **Relatedness within bioprojects.** In public repositories such as ENA, sequencing data are usually uploaded per individual projects, and ENA accession ranges often contain highly phylogenetically related genomes.
3. **Species clusters.** Individual microbial species form clusters in public repositories such as ENA ^35^.
4. **Sampling biases.** Public repositories exhibit prevalent sampling biases, enabling a rough classification of bacterial species into two categories: highly sampled and sparsely sampled (see, e.g., Fig. 1 in ref ^16^).

Altogether, this understanding led to the following general heuristic for batching genomes in comprehensive genome corpuses:

1. Cluster genomes based on their species. Specifically, identify the species of each genome, and then treat all genomes belonging to the same species as individual clusters.
2. Within each cluster, arrange genomes in the lexicographic order of their accessions, to maximize the chance that highly related genomes, sequenced at the same time, stay in the same batch in the subsequent steps,
3. Iterate over individual species clusters and compare their size with a predefined threshold (‘`batch-min-size’` in MiniPhy):

a. size≥threshold: Classify the species as highly sampled and proceed according to Step 5.
b. <threshold: Classify the species as sparsely sampled and proceed according to Step 4
4. Merge all sparsely sampled species clusters into a single pseudo-cluster called a dustbin, proceeding in the order of lexicographically sorted species names (while preserving the order of genomes within each cluster).
5. Split the dustbin pseudo-cluster into batches of a predefined size (‘`dustbin-batch-max-size’` in MiniPhy).
6. Split each highly sampled species cluster into batches of a predefined size (‘`batch-max-size`’ in MiniPhy).

The calibration of this heuristic was performed empirically, in the environment of the Harvard O2 cluster, with the paratemers adjusted based on observed performance. In particular, if the Mashtree/Attotree inference ^34^ or XZ compression of any batch exceeded a predefined time limit, the `batch-max-size` or `dustbin-batch-max-size` parameters were modified accordingly.

The resulting heuristic, including the default parameters, is provided in the MiniPhy repository in the ‘`create_batches.py’` script. The heuristic is also summarized, including the specific parameters used for 611k and the BIGSIdata, in **Supplementary Fig. 1**.

### Supplementary Note 4. Comparison of the Phylign and BLAST approaches

As tools for alignment against very large genome databases, Phylign and BLAST share many similarities, but at the same time, they differ in several key aspects. First, while Phylign is tailored specifically for bacterial genomes, BLAST is typically used with databases that encompass more types of sequences, including genes, transcripts, and genomes of non-bacterial organisms. Second, both tools produce alignments and compute alignment scores; however, while BLAST, computing local alignments, complements the score with an E-value to quantify the expected number of alignments of similar quality occurring by chance, Phylign targets longer alignments (primarily semiglobal, but can be adjusted by modifying Minimap parameters) and does not include E-values. Third, while both tools compute alignments using heuristic approaches, BLAST uses a seed-and-extend procedure, applied at the level of the entire database, whereas Phylign initially pre-filters target genomes using *k*-mer-based methods and then applies Minimap’s seed-chain-align procedure ^41^ at the level of individual reference genomes. In summary, Phylign and BLAST are designed for partially overlapping use cases, but they use different computational strategies.

## Supplementary Tables

**Supplementary Table 1:** Five test collections used for the calibration and evaluation of phylogenetic compression. Characteristics of the genome collection used for calibrating and evaluating phylogenetic compression throughout the paper. Within-genome *k*-mer duplicates refer to the proportion of *k*-mer occurrences (*k*=31, canonical *k*-mers) that disappear when transforming genome assemblies to their corresponding de Bruijn graphs; the fact that this proportion is always low for microbial genomes, even for complete assemblies, suggests that de Bruijn graphs are a faithful representation of microbial genomes and the *k*-mer content can be used for quantifying data redundancy.

| Collection | Description |  |  | Size |  | Diversity |  |  | Characteristics |  |  |  |
| --- | --- | --- | --- | --- | --- | --- | --- | --- | --- | --- | --- | --- |
| | Original samples | Genome representation | Data source | Nb. of genomes | Total sequence length | Nb. of species | Nb. of distinct $k$-mers | Within-genome $k$-mer duplicates | Unified construction pipeline | Data quality | Data volume | Repetitiveness |
| GISP | *N. gonorrhoeae* isolates | Draft assemblies<sup>70,79</sup> | .tar.gz file (726 MB)<br>https://doi.org/10.5281/zenodo.2618836 | 1,102 | 2.36 Gbp | 1 | 4.18 M | 2.02% | Yes | Very high | Low | Very high |
| NCTC3k | Bacterial isolates | Complete and draft assemblies<sup>96</sup> | .gff files (6.48 GB)<br>ftp://ftp.sanger.ac.uk/pub/project/pathogens/NCTC3000<br>Converted to FASTA and uploaded to:<br>https://doi.org/10.5281/zenodo.4838517<br>.fa.gz files (1.25 GB) | 1,065 | 4.35 Gbp | 259 | 992 M | 2.80% | Partially | High | Low | Medium |
| SC2 | SARS-CoV-2 isolates | Complete assemblies<sup>74</sup> | .xz file (201 MB)<br>http://gisaid.org | 590,779 | 17.6 Gbp | 1 | 1.85 M | 0.000700% | No | Low | High | Very high |
| 661k | Bacterial isolates | Draft assemblies<sup>97</sup> | .fa.gz files (805 GB)<br>http://ftp.ebi.ac.uk/pub/databases/ENA2018-bacteria-661k | 661,405 | 2.58 Tbp | 2,336 (est., ref<sup>97</sup>) | 44.3 G | 0.846% | Yes | Medium | Very high | High |
| BIGSIdata | Bacterial and viral isolates | de Bruijn graphs<sup>17</sup> | McCortex files (16.7 TB)<br>http://ftp.ebi.ac.uk/pub/software/bigsi/nat_biotech_2018/ctx | 425,160 | 1.68 Tbp<sup>a</sup> | 1,443 (est., ref<sup>17</sup>) | 41.1 G | - | Yes | Low | Very high | High |

**Footnotes:** <sup>a</sup> Computed as the total length of unitigs of the individual de Bruijn graphs.

**Supplementary Table 2:**
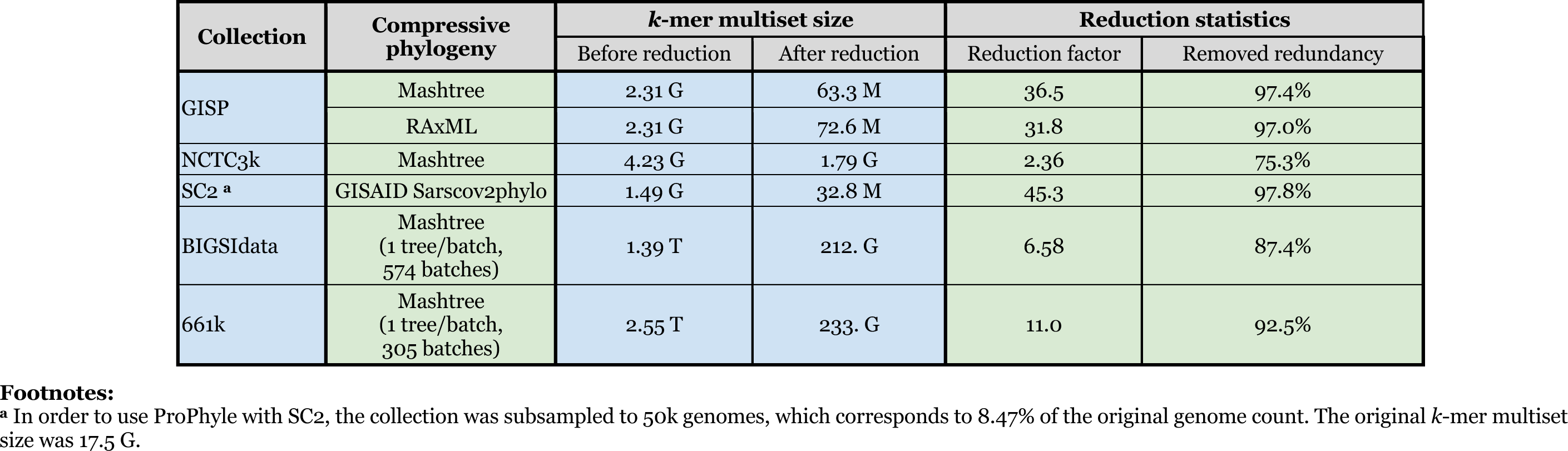
Proportion of redundancy explained by compressive phylogenies in the five test collections. The amount of reduction of genomic *k*-mer content (*k*=31, canonical *k*-mers) via *k*-mer propagation along compressive phylogenies. *k*-mer multisets correspond to the unions of *k*-mer sets before and after *k*-mer propagation, reduction factor is the ratio of their sizes, and removed redundancy quantifies the proportion of removed *k*-mers among the removable ones (100% if each *k*-mers was entirely associated with a single subtree). In the case of BIGSIdata and 661k, a phylogeny was built for each batch independently.

**Supplementary Table 3:**
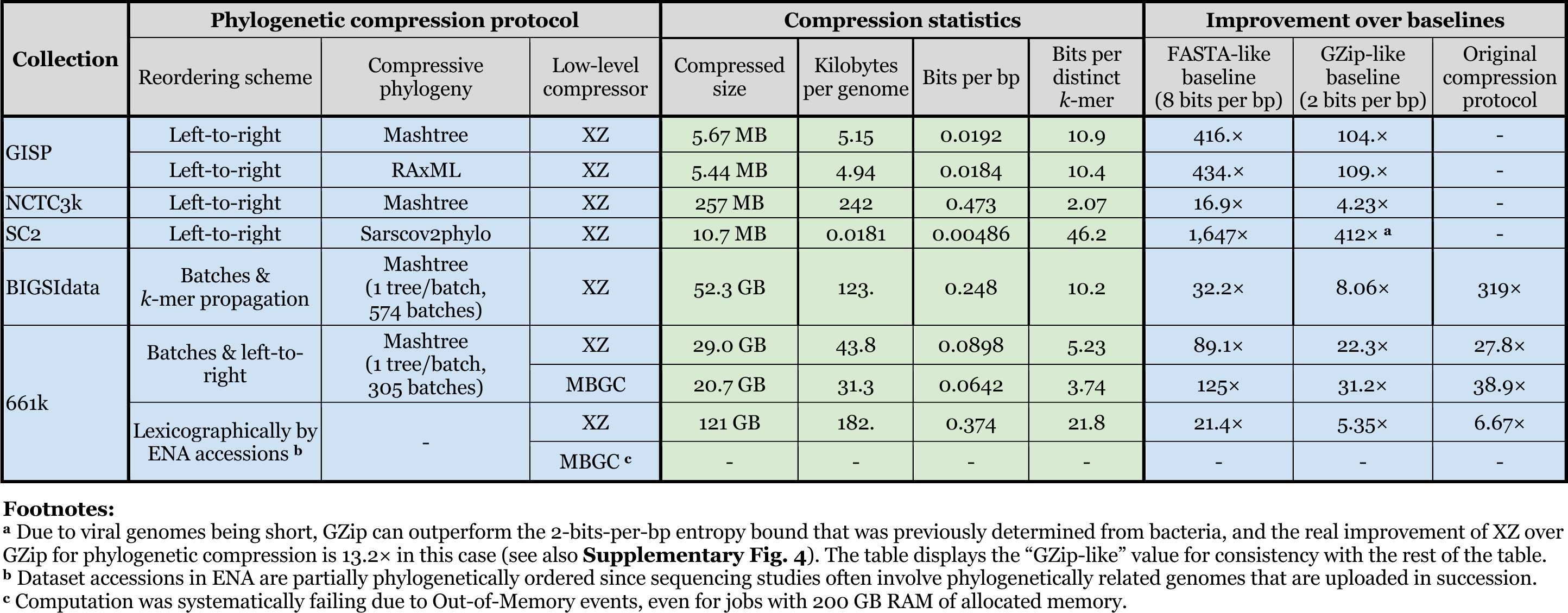
Results of phylogenetic compression. Size of the resulting files, mean space per single genome, bits per single base pair in the data, bits per distinct canonical *k*-mer (k=31), The three baselines include a FASTA-like baseline computed as 8 bits per single character (i.e., FASTA without sequence headers and EOLs), a GZip-like baseline (2 bits per bp), and the file size with original compression protocol.

| Collection | Phylogenetic compression protocol |  |  | Compression statistics |  |  |  | Improvement over baselines |  |  |
| --- | --- | --- | --- | --- | --- | --- | --- | --- | --- | --- |
| | Reordering scheme | Compressive phylogeny | Low-level compressor | Compressed size | Kilobytes per genome | Bits per bp | Bits per distinct $k$-mer | FASTA-like baseline (8 bits per bp) | GZip-like baseline (2 bits per bp) | Original compression protocol |
| GISP | Left-to-right | Mashtree | XZ | 5.67 MB | 5.15 | 0.0192 | 10.9 | 416.× | 104.× | - |
|  | Left-to-right | RAxML | XZ | 5.44 MB | 4.94 | 0.0184 | 10.4 | 434.× | 109.× | - |
| NCTC3k | Left-to-right | Mashtree | XZ | 257 MB | 242 | 0.473 | 2.07 | 16.9× | 4.23× | - |
| SC2 | Left-to-right | Sarscov2phylo | XZ | 10.7 MB | 0.0181 | 0.00486 | 46.2 | 1,647× | 412× | - |
| BIGSIdata | Batches & $k$-mer propagation | Mashtree (1 tree/batch, 574 batches) | XZ | 52.3 GB | 123. | 0.248 | 10.2 | 32.2× | 8.06× | 319× |
| 661k | Batches & left-to-right | Mashtree (1 tree/batch, 305 batches) | XZ | 29.0 GB | 43.8 | 0.0898 | 5.23 | 89.1× | 22.3× | 27.8× |
|  |  |  | MBGC | 20.7 GB | 31.3 | 0.0642 | 3.74 | 125× | 31.2× | 38.9× |
|  | Lexicographically by ENA accessions<sup>b</sup> | - | XZ | 121 GB | 182. | 0.374 | 21.8 | 21.4× | 5.35× | 6.67× |
|  |  |  | MBGC<sup>c</sup> | - | - | - | - | - | - | - |

**Supplementary Table 4:** Software and data provided for download. The table lists the developed software for phylogenetic compression and provides links to all phylogenetically compressed versions of the test collections (with the exceptions of SC2 that could not be published due to the licensing restrictions of GISAID).

| Type | Name | Description | URL |
| --- | --- | --- | --- |
| Software | Phylign | Snakemake pipeline | https://github.com/karel-brinda/phylign |
|  | MiniPhy | Snakemake pipeline | https://github.com/karel-brinda/miniphy |
|  | MiniPhy-COBS | Snakemake pipeline | https://github.com/leoisl/miniphy-cobs |
|  | De-MiniPhy-BIGSIdata | Client program to download and decompress de Bruijn graphs from the BIGSIdata collection | https://github.com/karel-brinda/de-miniphy-bigsidata |
|  | ProPhyle (modified, v0.3.3) | ProPhyle metagenomic classifier | https://github.com/prophyle/prophyle |
|  | COBS (modified, v0.3) | COBS *k*-mer indexer | https://github.com/iqbal-lab-org/cobs |
|  | Attotree | A fast reimplementaion of Mashtree functionality | https://github.com/karel-brinda/attotree |
| Phylogenetically compressed genome collections | NCTC3k | Assemblies (XZ) | https://doi.org/10.5281/zenodo.5533354 |
|  | BIGSIdata | De Bruijn graphs (simplitigs after *k*-mer propagation; XZ) | https://doi.org/10.5281/zenodo.5555253 |
|  | 661k | Assemblies (XZ) | https://doi.org/10.5281/zenodo.4602622 |
|  |  | Assemblies (MBGC) | https://doi.org/10.5281/zenodo.6347064 |
|  |  | *k*-mer index (COBS; XZ) | https://doi.org/10.5281/zenodo.7313926 |
|  |  |  | https://doi.org/10.5281/zenodo.7313942 |
|  |  |  | https://doi.org/10.5281/zenodo.7315499 |
|  | 661k-HQ | *k*-mer index (COBS; XZ) | https://doi.org/10.5281/zenodo.6849657<br>https://doi.org/10.5281/zenodo.6845083 |

**Supplementary Table 5:** Compressibility of different variants of BIGSI indexes for the 661k/661k-HQ collections.

| Collection | Protocol |  |  |  | Compression statistics |  |  |
| --- | --- | --- | --- | --- | --- | --- | --- |
|  | Index variant | Mode of construction | Low-level compression | Comment | Size | Total improvement over baseline | Improvement over uncompressed |
| 661k | COBS-compact <sup>a</sup> | Per entire collection | None <sup>b</sup> | Baseline (with adaptive Bloom filters) | 937. GB | 1.00× | 1.00× |
|  |  |  | XZ <sup>c</sup> | Direct compression | 243. GB | 3.86× | 3.86× |
|  | COBS-classic <sup>d</sup> | Per MiniPhy batch; columns sorted left-to-right | None | Reordered data, Bloom filter size fixed per batch | 2.46 TB | 0.380× | 1.00× |
|  |  |  | XZ | Phylogenetic compression | **110. GB** | **8.51×** | 22.5× |
| 661k-HQ | COBS-compact <sup>a</sup> | Per entire collection | None | Baseline (with adaptive Bloom filters) | 893. GB | 1.00× | 1.00× |
|  |  |  | XZ <sup>c</sup> | Direct compression | 205. GB | 4.35× | 4.35× |
|  | COBS-classic <sup>d</sup> | Per MiniPhy batch; columns sorted left-to-right | None | Reordered data, Bloom filter size fixed per batch | 1.06 TB | 0.842× | 1.00× |
|  |  |  | XZ | Phylogenetic compression | **72.8 GB** | **12.3×** | 14.5× |

<sup>a</sup> As the COBS-compact classifies datasets to be indexed into bins based on the number of *k*-mers, it is at the same time also grouping phylogenetically related genomes across the whole database into the same bins. The resulting file is thus moderately compressible using XZ, even though the resulting archive is not suitable for downstream applications because of its size and the associated overheads.
<sup>b</sup> Provided for download on http://ftp.ebi.ac.uk/pub/databases/ENA2018-bacteria-661k/.
<sup>c</sup> Streamed compression of COBS subindexes that are internally created by COBS based on the number of *k*-mers in individual datasets.
<sup>d</sup> Besides better compressibility by general compressors, COBS-classic brings an additional benefit of decreasing the associated false positive error rate for a majority of the datasets under indexing.

**Supplementary Table 6:** Disk space requirements of Phylign with the 661k-HQ collection. The requirements correspond to the version of the database as used by Phylign in **Supplementary Table 6**.

| Component | Size requirements<br>[GB] | Kilobytes per<br>genome <sup>a</sup> | Bits per bp <sup>a</sup> |
| --- | --- | --- | --- |
| Assemblies <sup>b</sup> | 29.2 | 45.6 | 0.0942 |
| COBS | 72.8 | 114. | 0.235 |
| **Total** | **102.** | **159.** | **0.329** |

**Supplementary Table 7:**
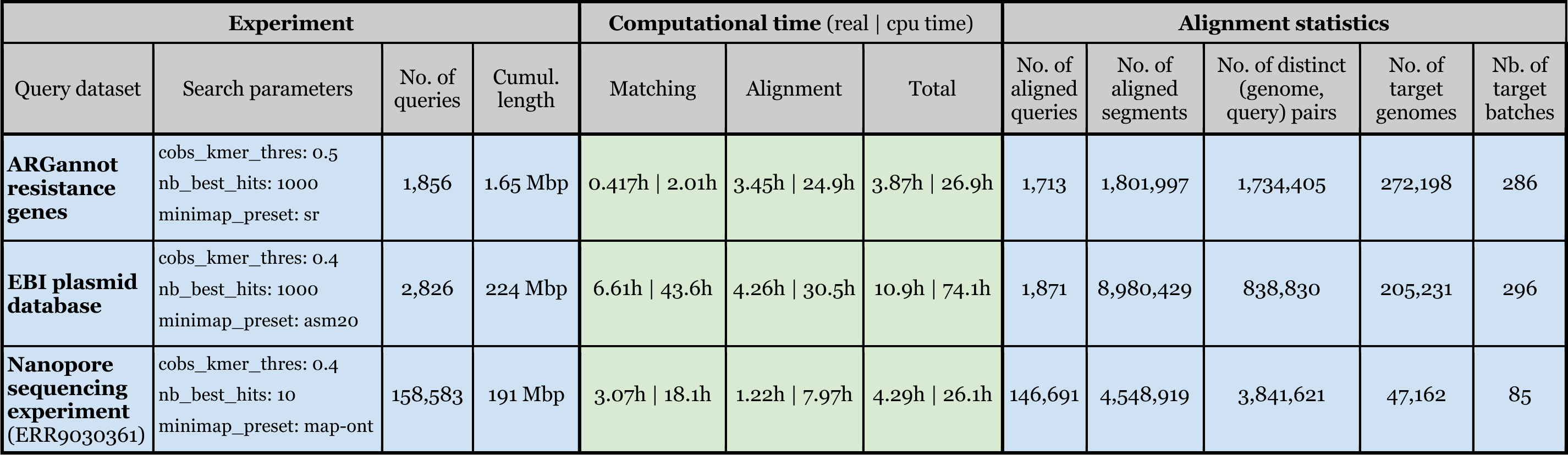
Results of BLAST-like search across the 661k-HQ collection on a desktop computer using Phylign. Timing and alignment results for resistance genes, EBI plasmids, and a nanopore sequencing experiment using Phylign, performed on an iMac with eight 4.2 GHz cores and 42.9 GB RAM (Methods). Search parameters were adjusted for the corresponding type of search based on the typical values in literature (Methods). All measurements were done with in-memory decompression (‘index_load_mode’ set to ‘mem-stream’) and maximal memory consumption set to 30 GB (the ‘max_ram_gb’ parameter). The resulting peak memory consumption was estimated to be 26.2 GB (Methods).

**Supplementary Table 8:** Reconstructed history of the BLAST NT database. The information about size of the BLAST nucleotide database (nt.gz) was retrieved from literature, its associated webpages, and other public repositories.

| Date | GZip size [GiB] | Length [Gbp] | Source <sup>a</sup> |
| --- | --- | --- | --- |
| 2002-01-01 | - | 7.372 | https://doi.org/10.1093/bioinformatics/btg250 (ref <sup>98</sup>) |
| 2003-02-01 | - | 8.33 | https://doi.org/10.1093/nar/gkh435 (ref <sup>99</sup>) |
| 2004-02-01 | - | 10 | https://doi.org/10.1093/nar/gkh435 (ref <sup>99</sup>) |
| 2007-07-01 | - | 21 | https://doi.org/10.1186/1471-2164-9-496 (ref <sup>100</sup>) |
| 2010-01-01 | - | 30 | https://doi.org/10.1186/1471-2105-11-340 (ref <sup>101</sup>) |
| 2012-10-11 | 10.7 | - | https://web.archive.org/web/20121011234515/http://ftp.ncbi.nih.gov/blast/db/FASTA |
| 2013-12-03 | 13.8 | - | https://web.archive.org/web/20201005113118/https://github.com/PathoScope/PathoScope/wiki/Building-Library |
| 2017-10-26 | 39.4 | - | https://doi.org/10.5281/zenodo.4382154 (ref <sup>102</sup>) |
| 2019-01-03 | 47 | - | https://openstack.cebitec.uni-bielefeld.de:8080/swift/v1/CAMI_2_DATABASES/ncbi_blast/nt.gz (a part of ref <sup>103</sup>) |
| 2020-04-05 | 67 | - | https://ftp.ncbi.nih.gov/blast/db/FASTA/ |
| 2020-07-05 | 72 | - | https://ftp.ncbi.nih.gov/blast/db/FASTA/ |
| 2020-08-04 | 77 | - | https://ftp.ncbi.nih.gov/blast/db/FASTA/ |
| 2020-10-11 | 81 | - | https://ftp.ncbi.nih.gov/blast/db/FASTA/ |
| 2021-01-15 | 90 | - | https://doi.org/10.17044/scilifelab.21070063.v1 (ref <sup>104</sup>) |
| 2021-03-21 | 103 | - | https://web.archive.org/web/20210322230129/https://ftp-trace.ncbi.nih.gov/blast/db/FASTA/ |
| 2021-03-28 | 104 | - | https://web.archive.org/web/20210402195739/https://ftp.ncbi.nih.gov/blast/db/FASTA/ |
| 2021-10-18 | 138 | - | https://web.archive.org/web/20211020093701/ftp://ftp.ncbi.nih.gov/blast/db/FASTA/ |
| 2021-11-01 | 139 | - | https://ftp.ncbi.nih.gov/blast/db/FASTA/ |
| 2021-12-13 | 146 | - | https://ftp.ncbi.nih.gov/blast/db/FASTA/ |
| 2022-01-18 | 152 | - | https://ftp.ncbi.nih.gov/blast/db/FASTA/ |
| 2022-02-28 | 161 | - | https://web.archive.org/web/20220307133636/https://ftp-trace.ncbi.nih.gov/blast/db/FASTA/ |
| 2022-03-21 | 166 | - | https://web.archive.org/web/20220326071216/https://ftp-trace.ncbi.nih.gov/blast/db/FASTA/ |
| 2022-06-06 | 180 | - | https://web.archive.org/web/20220609211512/https://ftp.ncbi.nih.gov/blast/db/FASTA/ |
| 2022-06-20 | 187 | 783.58 | https://ftp.ncbi.nih.gov/blast/db/FASTA/ |
| 2022-08-06 | 205 | - | https://ftp.ncbi.nih.gov/blast/db/FASTA/ |
| 2022-10-17 | 214 | - | https://ftp.ncbi.nih.gov/blast/db/FASTA/ |
| 2022-11-01 | 216 | - | https://ftp.ncbi.nih.gov/blast/db/FASTA/ |

**Footnotes:** <sup>a</sup> All webpages except https://ftp.ncbi.nih.gov/blast/db/FASTA/ were retrieved on 2022-11-02.

## Supplementary Figures

**Supplementary Fig. 1:**
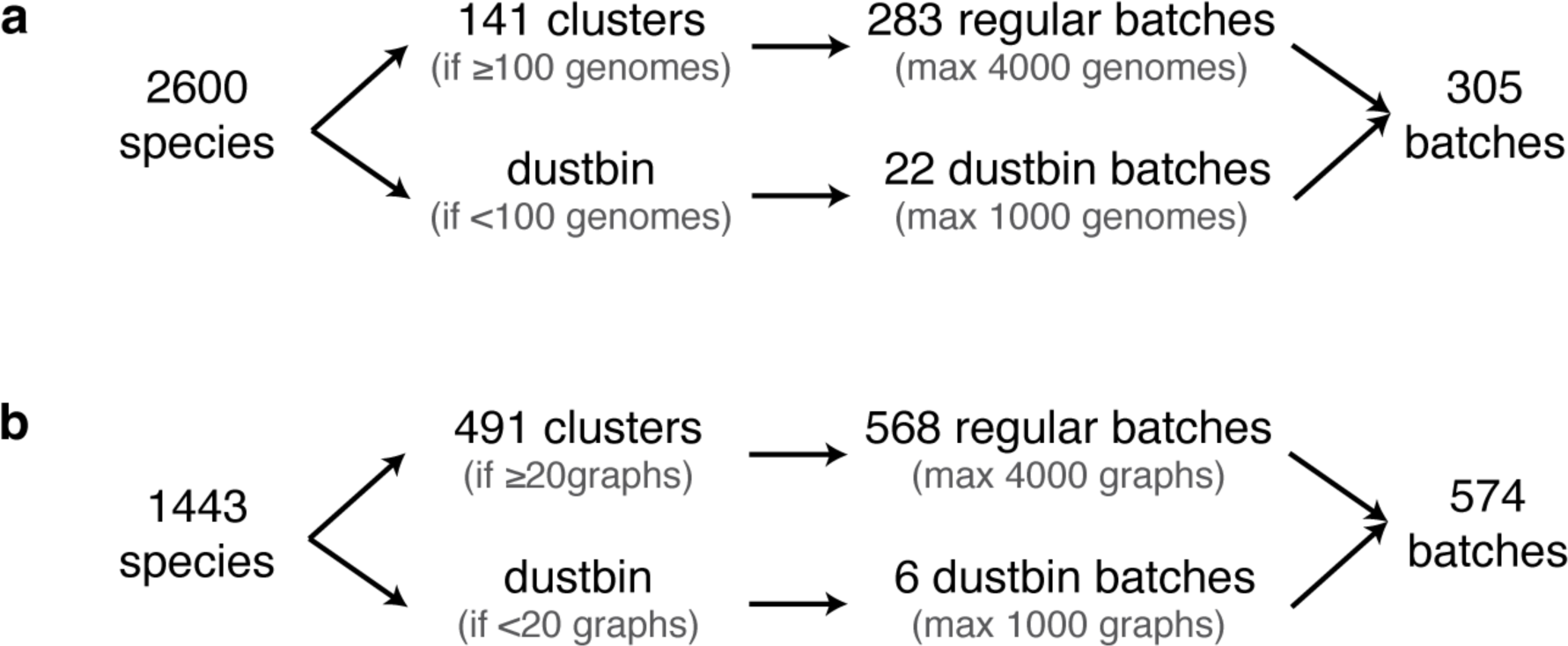
Batching strategies for the BIGSIdata and 661k collections. As a clustering strategy, genomes are grouped by individual species, and clusters that are too small are placed into a common pseudo-cluster called a dustbin. The obtained clusters and the dustbin are then divided into size-and diversity-balanced batches. The plot depicts the batching strategies used for the (**a**) 661k and (**b**) BIGSIdata collections. For further discussion of the batching, see **Supplementary Note 3**.

**Supplementary Fig. 2:**
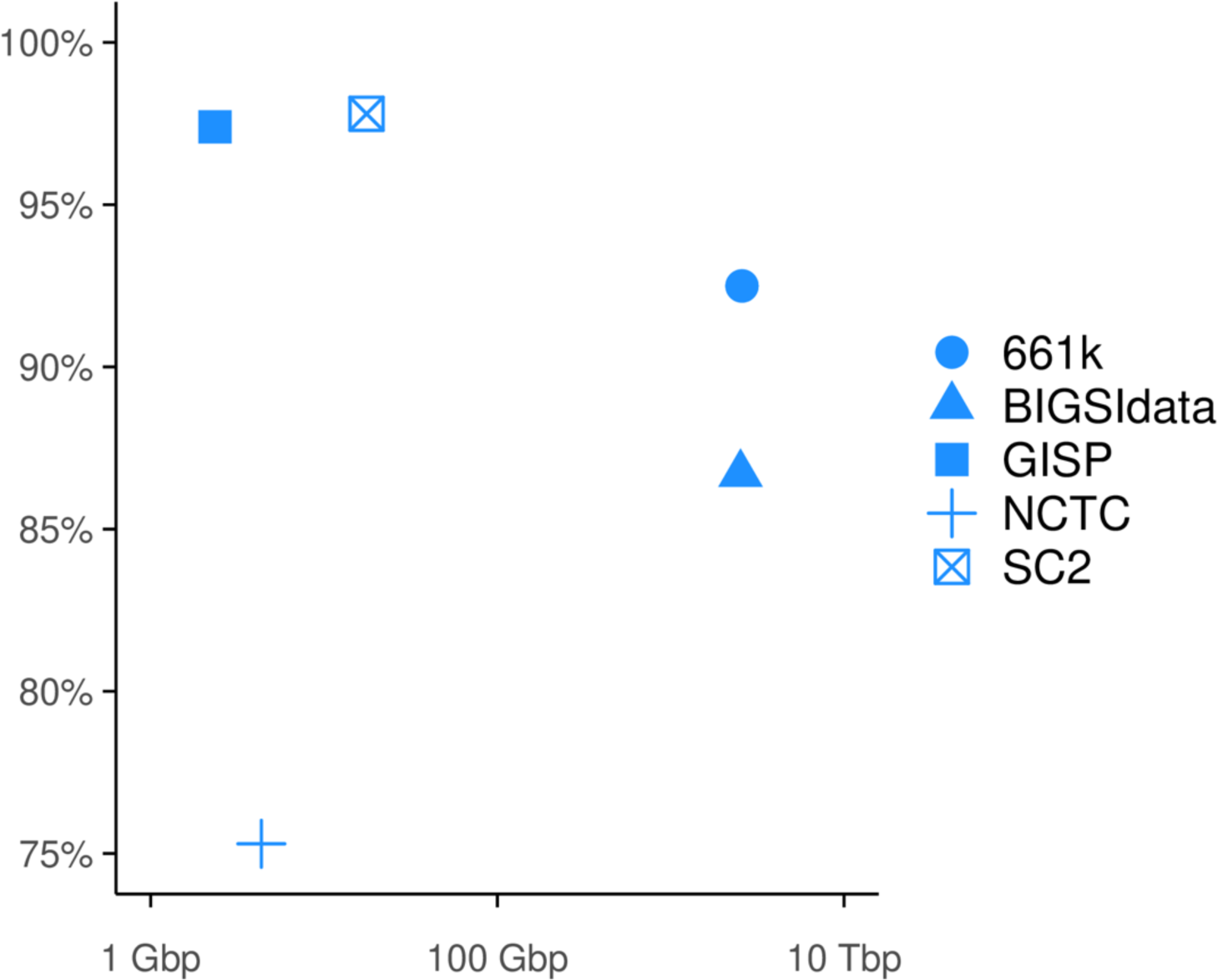
Quantification of phylogeny-explained data redundancy in the five test collections. The plot depicts the percentage of data redundancy that can be explained by the compressive phylogenies in each of the five test collections. The explained redundancy is measured by bottom-up *k*-mer propagation along the phylogenies performed by ProPhyle ^28,29^ and calculated as the proportion of *k*-mer duplicities removed by the propagation (see Methods for the formula). A *k*-mer distribution that is perfectly explained by the associated compressive phylogeny (i.e., all *k*-mers associated with complete subtrees) would result in 100% phylogeny-explained redundancy. The plot shows that for single-species batches (modeled by the GISP and SC2 collections), the majority of the signal can be explained by their compressive phylogenies, indicative of their extremely high phylogenetic compressibility. In contrast, high-diversity batches (modeled by the NCTC3k collection) have more irregularly distributed *k*-mer content due to horizontal gene transfer combined with sparse sampling, indicative of their lower compressibility (see **Supplementary Fig. 4**). Large and diverse collections, such as 661k and BIGSIdata, exhibit thus a medium level of phylogenetically explained redundancies, with the level depending on the amount of noise (higher for BIGSIdata and lower for 661k, as also visible in **Supplementary Fig. 7**).

**Supplementary Fig. 3:**
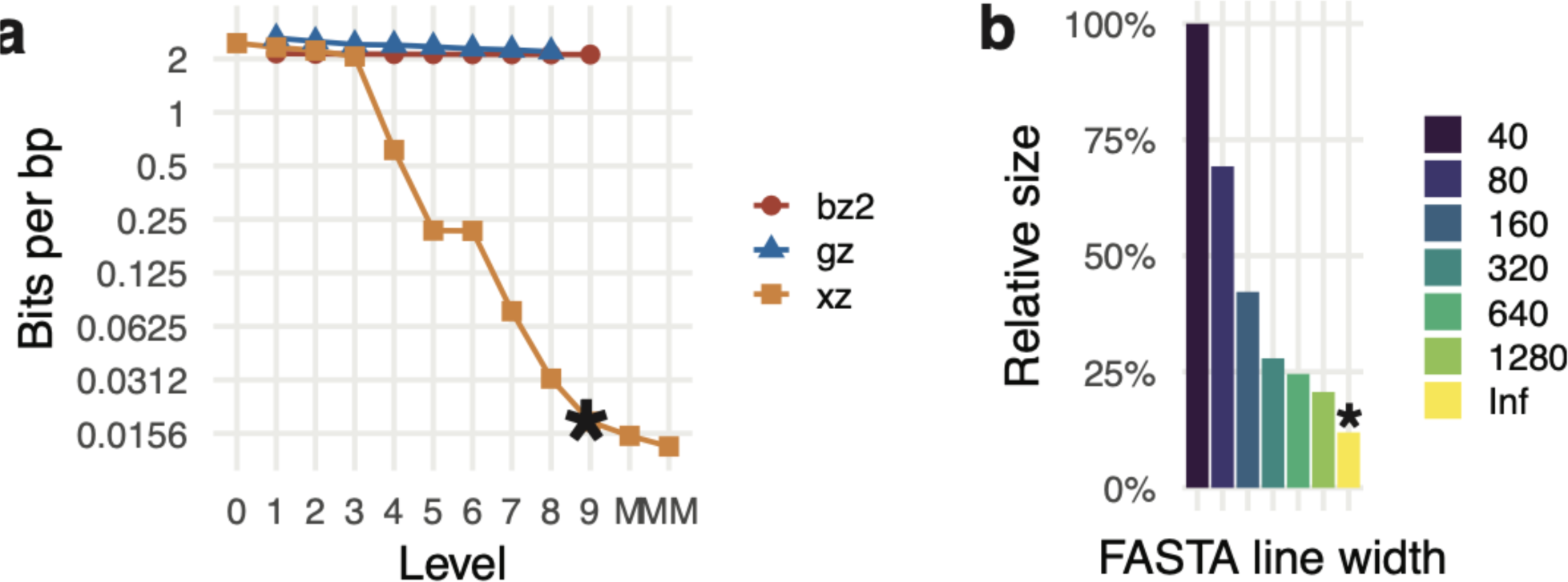
Calibration of XZ as a low-level tool for phylogenetic compression. The comparison was performed using the assemblies from the GISP collection, with genomes sorted left-to-right according to the Mashtree phylogeny. In both subplots, asterisk denotes the mode selected for phylogenetic compression in MiniPhy. **a)** The plot shows the compression performance XZ, GZip, and BZip2 in bits per base pair as a function compression presets (−1, -2, etc.) with single-line FASTA. Given the specific sizes of dictionaries and windows used in the individual algorithms and their individual presets, only XZ with a level ≥4 was capable of compressing bacterial genomes beyond the statistical entropy baseline (i.e., approximately 2 bits per bp). M and MM denote additional, manually tuned compression modes of XZ with an increased dictionary size (Methods), which slightly improved compression performance but at the same time substantially increased memory and CPU time and were thus not used in MiniPhy. **b)** The plot shows the impact of the FASTA line length on compression performance. With single-line FASTA (denoted by Inf), compression is improved to 12% of the 40 bps per line version. The plot highlights the importance of pre-formatting FASTA data before using general compressors such as XZ.

**Supplementary Fig. 4:**
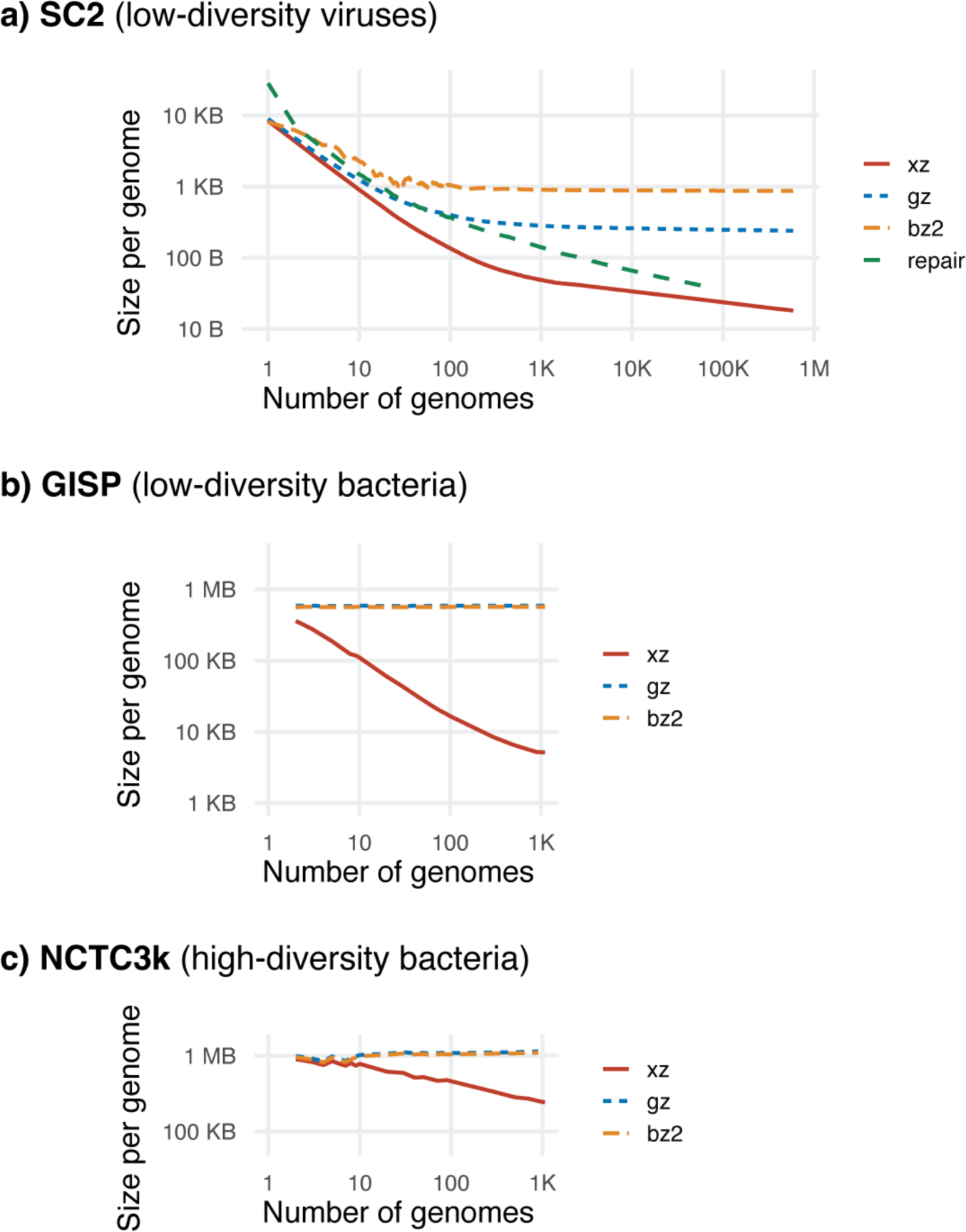
Comparison of three contrasting compression scaling modes of microbial collections. The plots compare on the scaling behavior of the XZ, GZip, BZip2, and RePair compressors on the SC2 (**a**), GISP (**b**), and NCTC3k (**c**) collections, depicting the space per single genome as a function of the number of jointly compressed genomes progressively increased on logarithmic scales. The results highlight several key findings. First, XZ consistently outperforms the other compressors. Second, for viral genomes all compressors are able to overcome the 2-bits-per-bp baseline thanks to their short genome length, but only XZ is able to compress beyond this limit for bacterial genomes (consistent with **Supplementary Fig. 3a**). Third, RePair compression can be nearly as effective as XZ for viruses, but its non-scalability limits its applicability to large datasets. Fourth, the compressibility of divergent bacteria is substantially limited even with the best compressors, with only a 4× improvement in per-genome compression for NCTC3k (while the highly compressible SC2 and GISP collections show 171× and 105× improvement for the same number of genomes).

**Supplementary Fig. 5:**
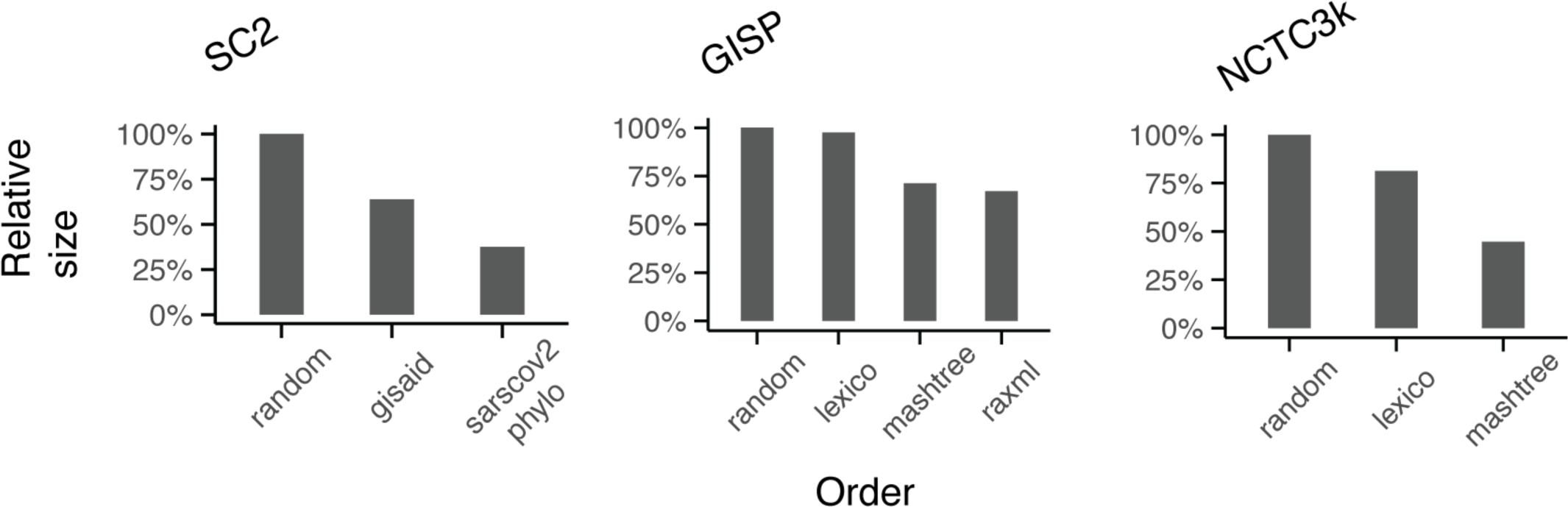
Impact of within-batch genome order on the compressibility of microbial collections. While a substantial part of the benefits of phylogenetic compression comes from the organization of genomes into batches of phylogenetically related genomes, proper genome reordering within individual batches is also crucial for maximizing data compressibility. The plots demonstrate that the impact of within-batch reordering grows with the amount of diversity included (GISP vs. NCTC3k) and with the number of genomes (GISP vs. SC 2). Accurate phylogenies inferred using RAxML provided only little benefits over trees computed using Mashtree (GISP).

**Supplementary Fig. 6:**
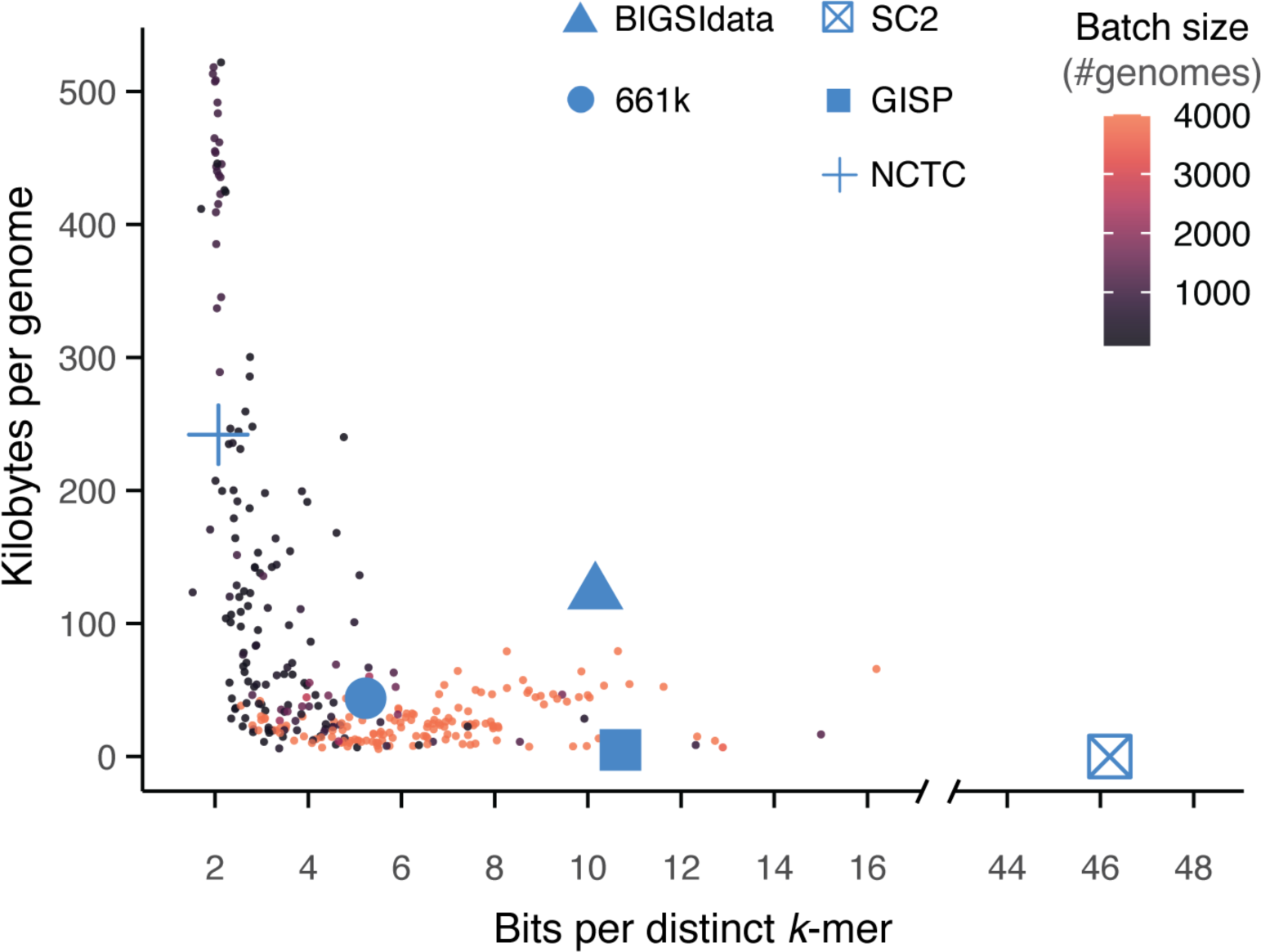
Compression tradeoffs for the five test collections and for individual batches of the 661k collection. The plot illustrates the tradeoff between the per-genome size after compression and the number of bits per distinct *k*-mer. The larger points represent individual genome collections and correspond to values from **Supplementary Table 3**. The smaller points represent individual batches of the 661k collection, with color indicating the number of genomes in each batch. Overall, the plot reveals the influence of genomic diversity on the resulting compression characteristics. The tradeoff follows an L-shaped pattern, where compression of genome groups with a high diversity leads to smaller space per *k*-mer but larger space per genome, and conversely for genome groups with a low diversity.

**Supplementary Fig. 7:**
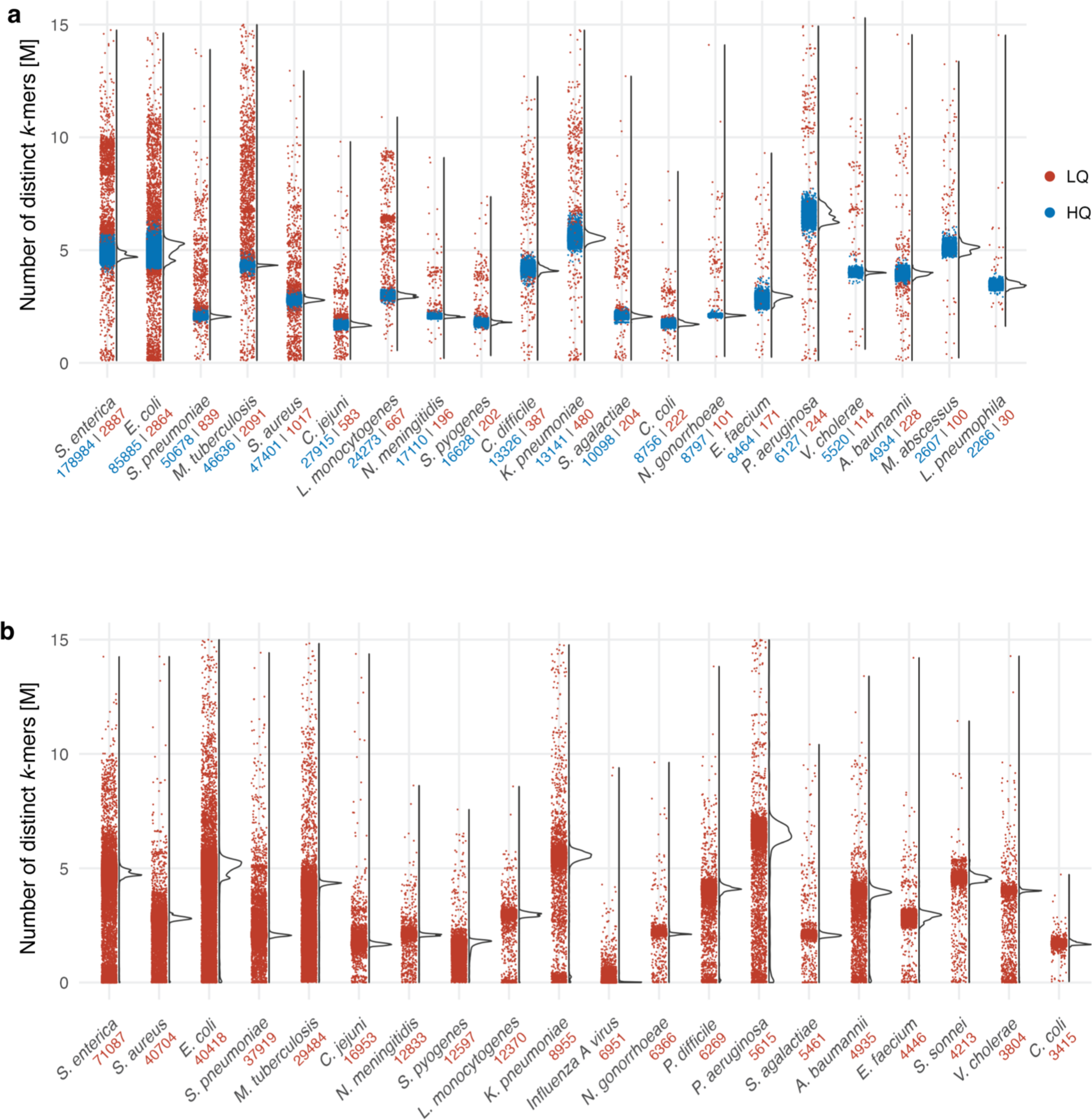
Distribution of the number of distinct *k*-mers in the top 20 species in (a) the 661k and (b) BIGSIdata collections. For the 661k collection, colors represent the quality of the assemblies (LQ: low-quality, HQ: high-quality), as determined as part of the quality control in ref ^97^. For BIGSIdata, no quality control information is available. The numbers below the species name indicate the number of samples within each category.

**Supplementary Fig. 8:**
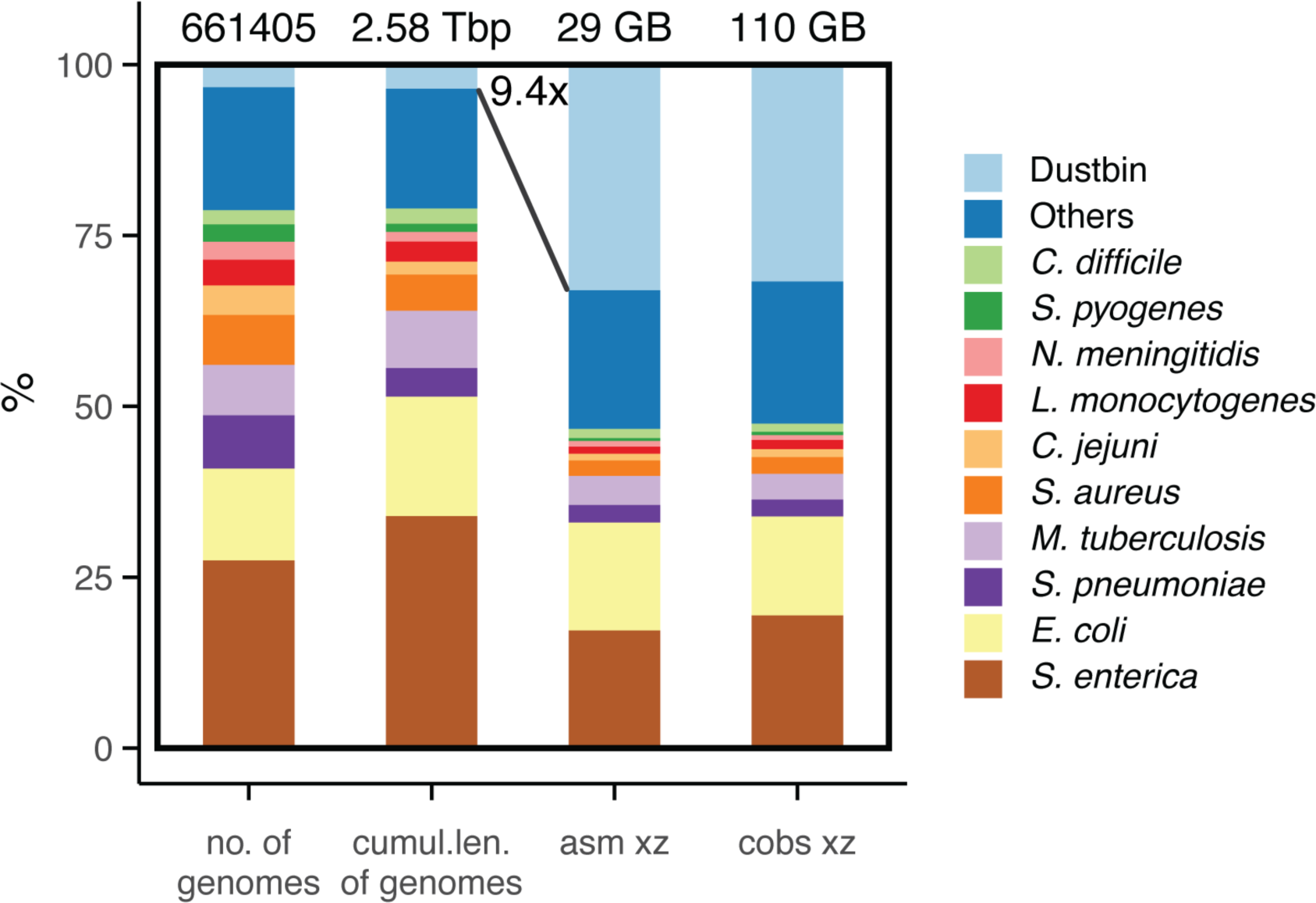
Proportions of top 10 species in the 661k collection before and after compression. The proportions of individual species (their corresponding batches) of the phylogenetically compressed 661k collection. The plot depicts the proportions of the top 10 species, the dustbin pseudo-cluster comprising divergent genomes, and the remaining species grouped in Others, while comparing the following four quantitative characteristics: the number of genomes, their cumulative length, the size of the phylogenetically compressed assemblies, and the size of the phylogenetically compressed COBS indexes. While transitioning from the number of genomes to their cumulative length has only a little impact on the proportions (only corresponding to different mean genome lengths of individual species), the divergent genomes occupy a substantially higher proportion of the collection after compression. Moreover, despite genome assemblies and *k*-mer COBS indexes are fundamentally different genome representations (horizontal vs. vertical, respectively), the observed post-compression proportions in them were nearly identical, indicative of that their compression is governed by the same rules.

**Supplementary Fig. 9:**
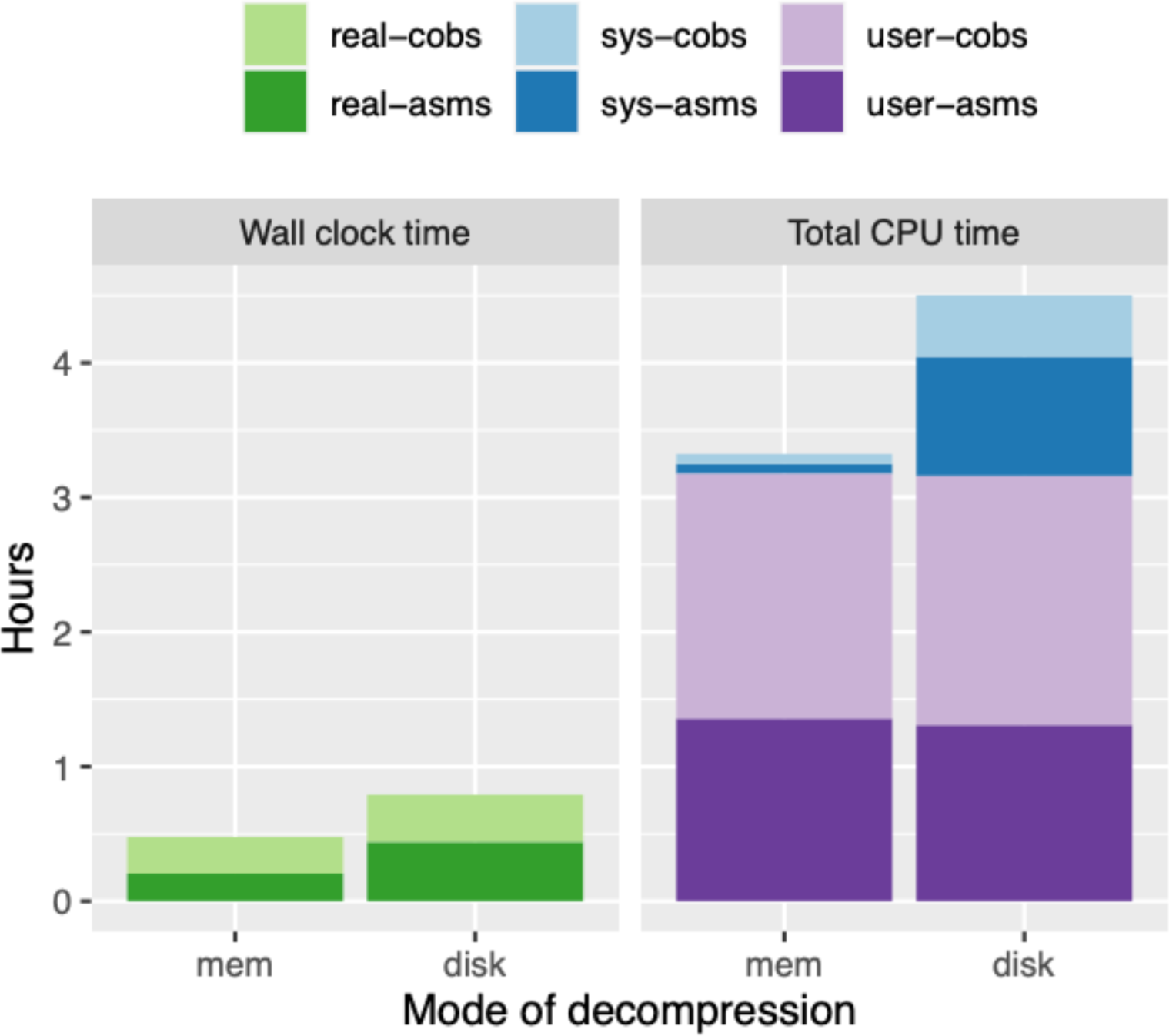
Time required for decompressing the Phylign database. The wall clock and total CPU time required to decompress the Phylign 661k-HQ database, both on a disk and in memory, measured on an iMac desktop computer with 4 physical (8 logical) cores. The decompression process in memory, which reflects the type of decompression used by Phylign, was completed under 30 mins, which is only a fraction of the typical duration of search experiments (see **Supplementary Tab. 6**).

## SUPPLEMENTARY FILES

Additional supplementary files are provided in a dedicated online repository on http://github.com/karel-brinda/phylogenetic-compression-supplement.

